# Machine Learning Elucidates Design Features of Plasmid DNA Lipid Nanoparticles for Cell Type-Preferential Transfection

**DOI:** 10.1101/2023.12.07.570602

**Authors:** Leonardo Cheng, Yining Zhu, Jingyao Ma, Ataes Aggarwal, Wu Han Toh, Charles Shin, Will Sangpachatanaruk, Gene Weng, Ramya Kumar, Hai-Quan Mao

## Abstract

For cell and gene therapies to become more broadly accessible, it is critical to develop and optimize non-viral cell type-preferential gene carriers such as lipid nanoparticles (LNPs).

Despite the effectiveness of high throughput screening (HTS) approaches in expediting LNP discovery, they are often costly, labor-intensive, and often do not provide actionable LNP design rules that focus screening efforts on the most relevant chemical and formulation parameters. Here we employed a machine learning (ML) workflow using well-curated plasmid DNA LNP transfection datasets across six cell types to maximize chemical insights from HTS studies and has achieved predictions with 5–9% error on average depending on cell type. By applying Shapley additive explanations to our ML models, we unveiled composition-function relationships dictating cell type-preferential LNP transfection efficiency. Notably, we identified consistent LNP composition parameters that enhance *in vitro* transfection efficiency across diverse cell types, such as ionizable to helper lipid ratios near 1:1 or 10:1 and the incorporation of cationic/zwitterionic helper lipids. In addition, several parameters were found to modulate cell type-preferentiality, including the ionizable and helper lipid total molar percentage, N/P ratio, cholesterol to PEGylated lipid ratio, and the chemical identity of the helper lipid. This study leverages HTS of compositionally diverse LNP libraries and ML analysis to understand the interactions between lipid components in LNP formulations; and offers fundamental insights that contribute to the establishment of unique sets of LNP compositions tailored for cell type-preferential transfection.

## Introduction

Gene and cell therapies have increasingly become a promising approach to combat a variety of diseases from genetic disorders to cancer^1^. Although these therapeutics have proven efficacious across a variety of applications, reliable and targeted delivery of genetic payloads to specified *in vivo* and *in vitro* cell and tissue targets remains elusive^2^. Most FDA-approved gene and cell therapies rely on viral gene delivery, which are limited by payload packaging capacity, compromised therapeutic outcomes due to pre-existing antibodies, and patient immune response, resulting in efficacy and safety concerns^3–5^. These challenges continue to provide impetus for the development of non-viral carriers such as lipid or polymeric nanoparticles as alternative solutions for gene therapies. Lipid nanoparticles (LNPs) have demonstrated remarkable potential as an effective carrier in nucleic acid delivery *in vitro* and *in vivo*^6–8^. The approvals of siRNA LNPs (Onpattro) and two mRNA LNP-based SARS-COV-2 vaccines, Spikevax® (Moderna) and Comirnaty® (BioNTech/Pfizer), have solidified LNPs as a leading non-viral carrier for gene therapeutics^9^. Despite success in gene vaccine delivery, LNPs are not a “one formulation fits all” delivery modality; and the design space of LNP formulations must be expanded and refined to address factors such as cell type-preferential transfection, tissue/organ targeting, and route of administration^10^.

LNPs are classically composed of four lipid components: ionizable cationic lipids, helper lipids, cholesterol, and PEGylated lipids^10,11^. Ionizable cationic lipids, which typically incorporate tertiary amines with pH-sensitive protonation equilibria, primarily package anionic payloads (nucleic acids) and facilitate endosomal escape. Several studies have highlighted the importance of incorporating additional lipids such as helper lipids to promote cellular association and targeting, cholesterol for fusogenicity, structural integrity, and stability, and PEGylated lipids to modulate plasma protein interactions, LNP colloidal stability, and LNP biodistribution and pharmacokinetics *in vivo*^10,12–14^. Nevertheless, while the roles of ionizable lipids have been extensively mapped, the roles of other lipid components, particularly helper lipids, and the combined interactions of lipid components with each other remains poorly understood, thus limiting our ability to rationally design and screen LNP compositions. LNP optimization is notoriously inefficient because LNP composition parameters compose a vast and relatively unexplored design space. Unless LNP optimization is guided and informed by fundamental insights into the roles of lipid chemistry and formulation parameters, LNP design will largely remain subject to trial-and-error experimentation. Therefore, prudent strategies to probe and characterize this vast design space are necessary.

High throughput screening (HTS) studies have become an increasingly routine approach to optimize new LNP formulations. The majority of LNP HTS studies have focused on synthesizing and screening new ionizable lipids to achieve optimal LNP formulations. However, they offer very limited chemical insights into the roles of helper lipid chemical identity and overall formulation compositions^15–18^. Our recent study on HTS of LNPs highlighted that designing effective LNP formulations requires extensive optimization of composition and careful selection of compatible helper lipid chemistries^19^. Specifically, our HTS identified top LNP compositions from a 1080-formulation library (varying helper lipid charge and lipid component ratios) that effectively transfected hepatocarcinoma cells *in vitro* and more importantly achieved successful delivery to the liver after intrahepatic or intravenous injection. Another recent study showed that cell specific transfection can be achieved by modulating formulation composition and helper lipid chemical identity without altering the ionizable lipid structure^20^.

Machine learning (ML) algorithms combined with downstream analysis techniques are promising to explore and understand the interplay between multiple design parameters governing LNP transfection efficiency. Developing and validating ML workflows that are robust to experimental noise can circumvent the reliance on exhaustive and tedious large-scale screening. Given sufficiently large well-curated training data, ML algorithms can identify complex relationships between input parameters (LNP features) and output responses (transfection performance). Several recent reports showcased the ability of ML approaches to predict how carrier attributes (*e*.*g*., size, polydispersity index, and _ζ_-potential) influence biological responses such as transfection efficiency and cell viability of LNPs and polyplexes^21–25^. ML also assisted the development of new ionizable lipid chemistries but the role of helper lipid chemistry and lipid component ratios has not been extensively explored within a ML framework^22,24^.

Several common ML algorithms have been utilized in the analysis of non-viral gene carrier systems. Most are considered complex “black box models”, which renders interpretation and elucidation of structure-function (transfection efficiency) relationships challenging. Shapley Additive Explanations (SHAP), an analysis framework grounded in game theory, has been shown to robustly interpret findings from ML analysis by providing a unified measure of feature importance^26^. Here, we used transfection dataset of an LNP library comprising 1,080 LNP formulations by varying LNP compositions and helper lipid chemistry. This compositionally diverse LNP library was tested in 6 cell types and their *in vitro* transfection performance assessed^19,20^. We developed a computational pipeline that will select, train, and refine high performing ML algorithms that predict transfection outcomes as a function of LNP design features in each cell type. We then conducted comparative SHAP analysis of feature importance to elucidate structure-function relationships governing cell type-preferential transfection. The integration of experimenter-directed HTS with a robust ML pipeline presents a new approach to confidently decipher the structure-function relationship of LNP formulations.

## Results and Discussion

### Helper lipid and LNP composition modulate cell type-preferential transfection efficiency

The dataset used to train the ML model was produced using an HTS approach for a nucleic acid-loaded LNP library that we generated and characterized previously^19,20,27^ (**Figure 1a**) in B16F10 (murine melanoma), HepG2 (human hepatocellular carcinoma), HEK293 (human embryonic kidney), PC3 (human prostatic adenocarcinoma), ARPE-19 (human retinal pigment epithelia), and N2a (murine neuroblast) cell lines. Each formulation composition contains DLin-MC3 as the ionizable lipid, cholesterol, DMG-PEG2000, and one helper lipid selected from a group of 6 carrying different charge types: cationic head groups (DOTAP, DDAB), zwitterionic head groups (DOPE, DSPC), and anionic head groups (18PG, 14PA) based on our previous screening studies^19^. The following four independent composition parameters were adjusted: (i) total molar percentage of ionizable lipid and helper lipid ranging from 20–80% (“DLin + helper lipid total %”), (ii) ionizable lipid to helper lipid molar ratio ranging from 1:200 (“DLin/helper lipid ratio”), (iii) protonatable nitrogen to phosphorous ratio (”N/P ratio”) ranging from 4 to 12, and (iv) cholesterol to PEGylated lipid molar ratio ranging from 1 to 500 (“Chol/PEG ratio”) (**Supplementary Table 1 and Supplemental Figure 1a**). By varying these composition parameters, 180 unique LNP formulations were generated from each helper lipid. Initially seven molecular descriptors were generated based on the helper lipid chemical structure using the most distinctive chemical structural features to parameterize the structure of the helper lipids (**Supplemental Table 2 and Supplemental Figure 1b**), and prioritized based on the chemical interpretability of the results^28^. For example, the number of positively/negatively charged centers and the number of hydrogen bond acceptors/donors quantify the capacity of helper lipids to form electrostatic interactions and hydrogen bonds within a pDNA LNP. Other features such as the number of double bonds, the total number of carbons atoms within the lipid tails, and calculated LogP (cLogP) may characterize the structural rigidity and lipophilicity of the helper lipid^11,29^. In addition to helper lipid chemical features, particle-level physiochemical properties such as LNP size and polydispersity, and _ζ_-potential were included (**Supplemental Figure 1c**).

**Table 1.** Model selection results for LNP transfection in B16F10 cells.

| **Model Type** | **LGBM** | **RF** | **XGB** | **DT** | **kNN** | **MLR** | **PLS** | **lasso** |
| --- | --- | --- | --- | --- | --- | --- | --- | --- |
| Mean Absolute Error | 0.055 | 0.059 | 0.060 | 0.070 | 0.103 | 0.103 | 0.104 | 0.106 |
| Spearman's Rank Correlation | 0.90 | 0.88 | 0.87 | 0.81 | 0.61 | 0.67 | 0.67 | 0.66 |
| Pearson's Correlation | 0.91 | 0.89 | 0.89 | 0.83 | 0.65 | 0.68 | 0.68 | 0.66 |

**Table 2.** Model refinement results using LGBM in different cell types.

| **Cell type** | **B16** | **HepG2** | **PC3** | **HEK293** | **N2a** | **ARPE-19** |
| --- | --- | --- | --- | --- | --- | --- |
| Number of features needed | 10 | 9 | 7 | 9 | 7 | 7 |
| Composition features retained | All | All | All | All | All | All |
| Helper lipid features retained (+) |  |  |  |  |  |  |
| Positively charged centers | + | + | + | + | + | + |
| cLogP | + | + | + | + | + | + |
| hydrogen bond donors | + | + | - | + | - | - |
| hydrogen bond acceptors | + | + | - | + | + | + |
| carbons in tails | + | + | - | - | - | - |
| double bonds in tails | + | - | + | + | - | - |
| Physicochemical features retained | None | None | None | None | None | None |
| MAE average | 0.049 | 0.049 | 0.051 | 0.083 | 0.058 | 0.093 |
| Pearson average | 0.91 | 0.70 | 0.67 | 0.85 | 0.90 | 0.88 |
| Spearman average | 0.93 | 0.88 | 0.90 | 0.83 | 0.92 | 0.89 |

**Figure 1.**
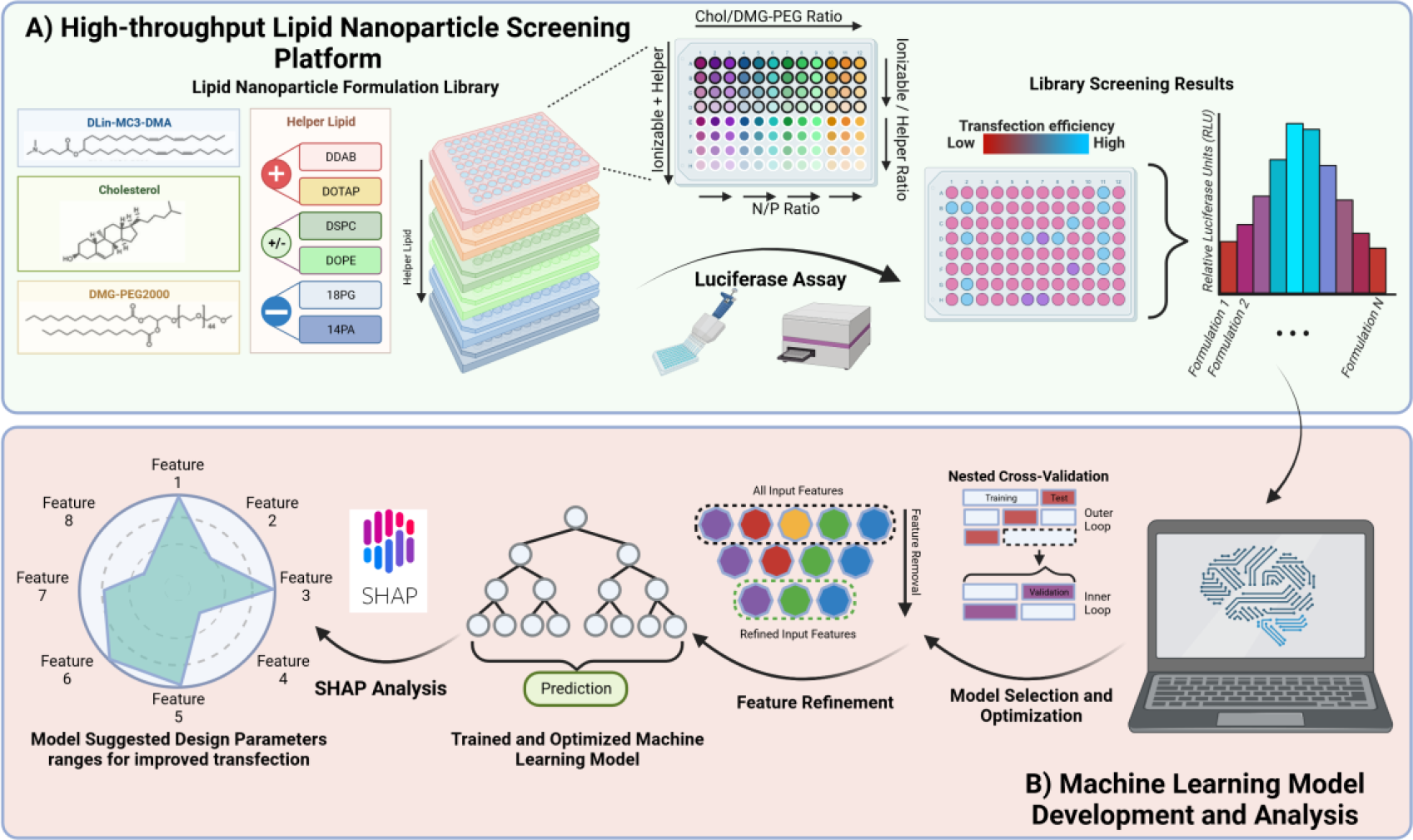
Schematic of high throughput screening and ML pipeline. **(a)** Data generation using an LNP library and screening platform tested in 6 different cell types. **(b)** Model selection and hyperparameter optimization using a nested cross validation approach and model Input feature refinement. Shapley additive explanations (SHAP) was used to analyze optimized models and elucidate design rules for cell type-preferential LNP formulations.

Transfection efficiency measured by the relative luminescence units (RLU) obtained with this LNP library was sufficiently broad (e^2^ - e^12^ RLU/well) for most cell types. Broadly, the transfection efficiency was sensitive to the cell type, the chemical identity of the helper lipid, and LNP composition (**Figure 2a**). We observed non-overlapping transfection efficiency trends in each cell type, indicated by the low (r < 0.35) or moderate rank correlation (0.36 < r < 0.67)^30^ of formulation-dependence in transfection efficiency between each pair of cell types (**Figure 2b**). A low Spearman’s rank correlation coefficient indicates that LNP features responsible for transfection efficiency diverge between cell types whereas a high Spearman’s rank coefficient suggests overlapping LNP design criteria shared by two cell types. For most part, LNPs that performed well in one cell type typically underperformed in others. The two exceptions were N2a and ARPE-19 cells, which were the most highly correlated cell lines with a Spearman’s rank correlation coefficient of 0.71. We attributed the high correlation to the shared neuronal origins of ARPE-19 and N2a cells^31,32^. Our results suggest that it may be possible to exploit interactions between LNP design features and certain cellular features of the target cell type to develop tailored LNPs for cell type-preferential transfection through modulation of the formulation compositions and helper lipid identity. Further analysis of the Spearman’s rank coefficients between subsets of formulations containing the same helper lipid revealed that specific formulation composition parameters were highly conserved while using DDAB, moderately conserved using DOTAP and DOPE, and weakly conserved in DSPC, 18PG, and 14PA. Our analysis establishes that the choice of helper lipid impacts the extent to which LNPs selectively mediate efficient pDNA delivery in one cell type but not in others (**Supplementary Figure 2**).

**Figure 2.**
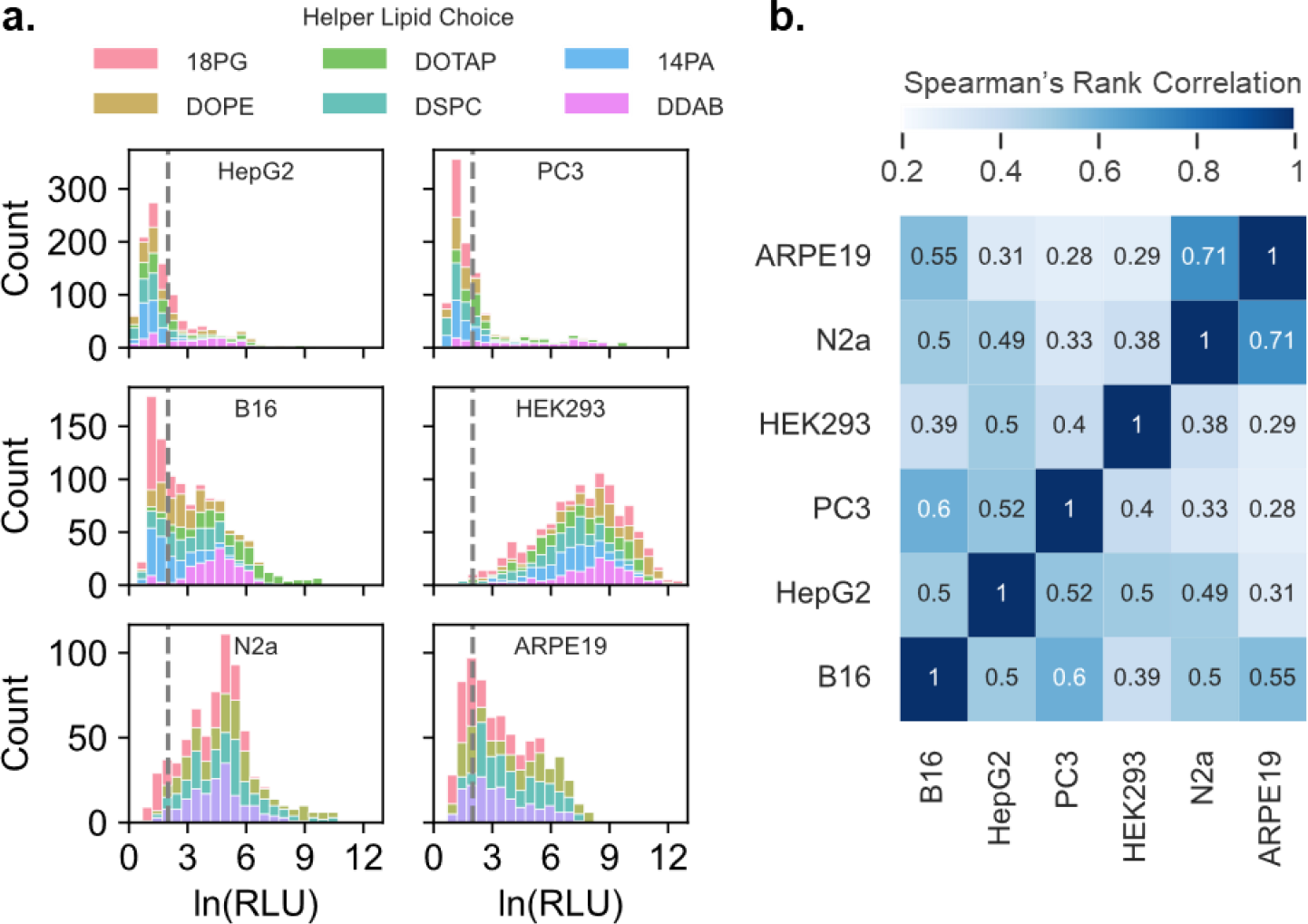
Preliminary Comparison of Screening Results across the cell lines. **(a)** Distribution of transfection efficiency of each cell type colored by helper lipid used in formulation. Gray dotted line indicates the value used to floor the dataset during preprocessing **(b)** Heatmap of Spearman’s rank correlation coefficient of LNP formulation transfection between cell types after data preprocessing.

### Decision tree-based algorithms describe complex non-linear relationships between LNP formulation features and transfection efficiency

The dataset was preprocessed to facilitate comparative analysis between cell type-preferential models (see Methods for details). Importantly, the transfection efficiency values within a specific cell type dataset were normalized between a range of 0 (min) to 1 (max) based on the maximum transfection achieved to eliminate the impact of overall cell type “transfectability” from downstream analysis.

Various ML algorithms are available to capture trends in datasets; however, certain algorithms may perform better than others depending on the complexity and type of relationship between the input and output parameters. Furthermore, while a model may be able to fit a relationship to training data precisely, researchers must be wary of the harmful impacts of over/underfitting and lack of training data influencing model robustness. To address these concerns, model selection was conducted to select the best performing algorithms with minimal overfitting and validate whether the training data was sufficiently large. A nested-cross validation approach was used to evaluate the performance and select the best performing models from a panel of regression analyses, nearest neighbors’ algorithms, and decision tree-based algorithms^33^. We used a 5-fold cross-validation data splitting strategy (outer loop) to randomly selected 80% of the dataset for model development and hyperparameter optimization (inner loop) and used the remaining 20% of the training dataset as a hold-out test set (**Figure 1b**). Within the inner loop, we applied another 4-fold cross-validation split and conducted 100 iterations of hyperparameter optimization using a random grid search approach to identify the best model hyperparameters (**Supplementary Table 3)**. The best model hyperparameters were selected by minimizing the “validation error” or the mean absolute error (MAE) between the inner loop validation set and model predictions. After selecting the best model hyperparameters, the “test error” was evaluated by calculating the MAE of model predictions on the outer loop hold-out data set. This process was repeated five times in total to identify five sets of optimized hyperparameters. Due to the rigorous inner loop model selection process, we identified many high performing models, so the final model was selected to reduce model overfitting by minimizing the difference between the validation and test error. Our robust model selection approach integrated rigorous hyperparameter optimization and balanced the validation and test error, thus ensuring that high performing and minimally overfitting model architectures were chosen.

**Table 3.**
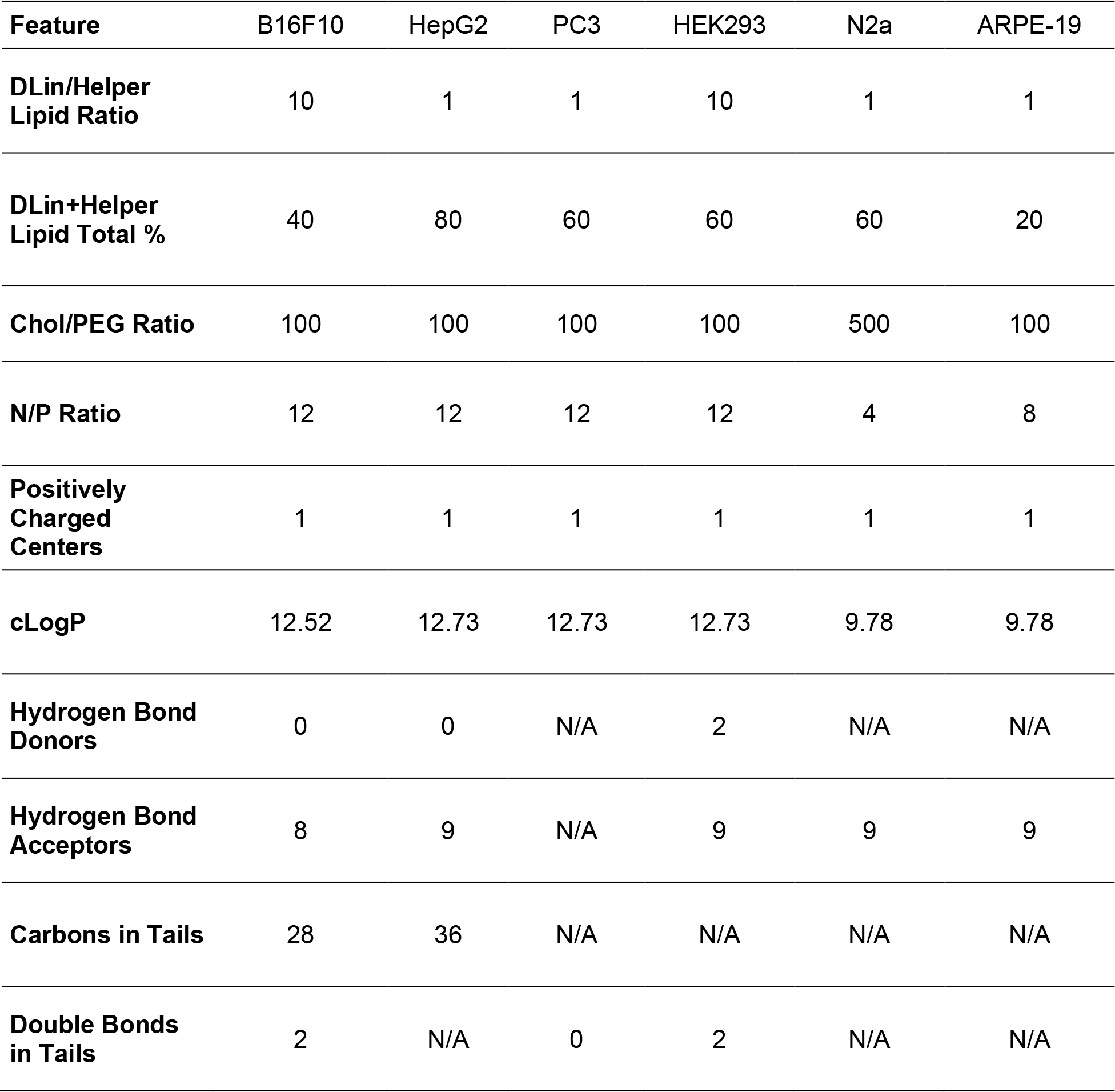
Machine learning elucidated design rules for optimal cell type transfection.

| **Feature** | **B16F10** | **HepG2** | **PC3** | **HEK293** | **N2a** | **ARPE-19** |
| --- | --- | --- | --- | --- | --- | --- |
| **DLin/Helper Lipid Ratio** | 10 | 1 | 1 | 10 | 1 | 1 |
| **DLin+Helper Lipid Total %** | 40 | 80 | 60 | 60 | 60 | 20 |
| **Chol/PEG Ratio** | 100 | 100 | 100 | 100 | 500 | 100 |
| **N/P Ratio** | 12 | 12 | 12 | 12 | 4 | 8 |
| **Positively Charged Centers** | 1 | 1 | 1 | 1 | 1 | 1 |
| **cLogP** | 12.52 | 12.73 | 12.73 | 12.73 | 9.78 | 9.78 |
| **Hydrogen Bond Donors** | 0 | 0 | N/A | 2 | N/A | N/A |
| **Hydrogen Bond Acceptors** | 8 | 9 | N/A | 9 | 9 | 9 |
| **Carbons in Tails** | 28 | 36 | N/A | N/A | N/A | N/A |
| **Double Bonds in Tails** | 2 | N/A | 0 | 2 | N/A | N/A |

Across all cell types, the best performing models were achieved for the B16F10, HepG2, and PC3 cell types using the LightGBM (LGBM) model achieving a low mean absolute error (MAE) of below 0.06 (*i*.*e*., predicted transfection efficiency with below 6% error on average) (**Supplementary Table 4**) when evaluated using a 5-fold cross-validation split. For the B16F10 cell type, model selection established that the LGBM model performed the best among all tested models (**Figure 3a**). The LGBM model achieved impressive Pearson’s correlations (r = 0.91), and Spearman’s rank (r = 0.90) on the outer loop hold-out datasets (**Figure 3b)**. Further *in vitro* validation emphasizes our strong model predictive abilities in unexplored feature spaces (**Supplementary Figure 3, Supplementary Table 4**). Interestingly, decision tree-based models (LGBM, XGB, RF, DT) performed significantly better than the other tested models, however, there was little difference in performance among the four decision tree-based models (**Figure 3a**). Unlike regression models, decision tree-based models fit data against ordered sets of yes/no decisions (“nodes”) to determine the predicted output. As such, this method fits non-linear, complex relationships more accurately than other algorithms such as linear regression-based, which are better suited for simpler, linear relationships described by explicit mathematical equations^34^. Given that the decision-tree based models most effectively analyzed the dataset, it suggests that the relationships between formulation parameters and transfection efficiency are non-linear and complex otherwise regression models would have performed equally well. Similar results have been reported in other comparable tasks where decision-tree based models outperform various other model architectures^35–40^. We achieved minimal error (MAE < 0.11), strong Pearson’s correlation (r > 0.8), and moderate to strong Spearman’s rank correlation (r > 0.6) between the predictions and hold-out experimental values across all cell types (**Supplementary Table 4, Supplementary Figure 4**). We also observed similar trends where decision-tree based models performed significantly better than regression and more specifically, we found LGBM models performed the best for all cell types (**Supplementary Figure 5**). Given the difficulty of modeling non-linear complex relationships between the formulation features and transfection across all cell lines, our results establish that decision-tree based ML models most reliably capture and model critical features underlying LNP transfection performance.

**Figure 3.**
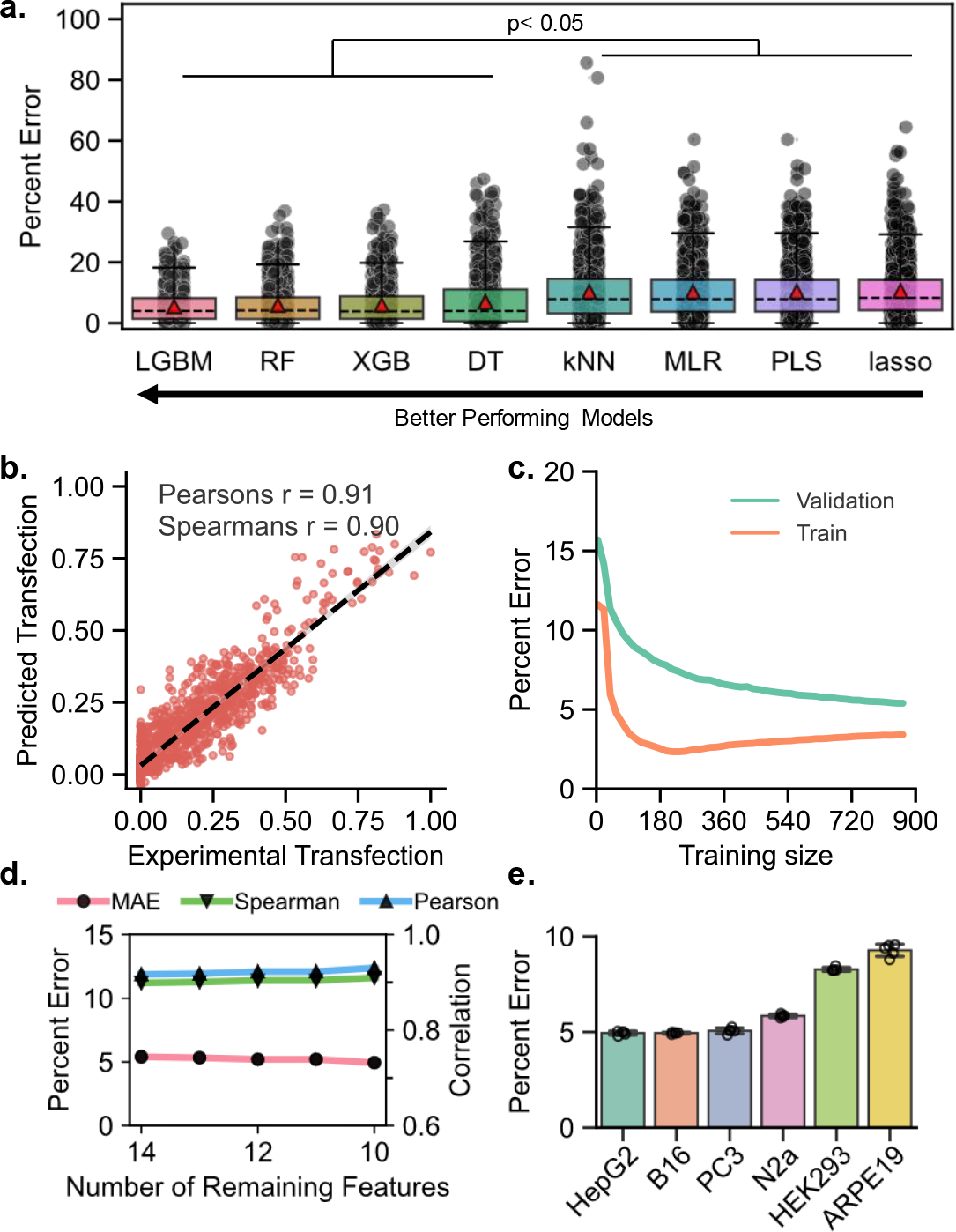
Model selection and hyperparameter optimization for B16F10 cell type. **(a)** Boxplot of hold-out set MAE aggregated across all 5-fold cross validation outer loop folds. Boxplot is centered around the MAE (red triangles), and dots represent individual hold-out set AE. Models are ranked from lowest (left) to highest (right) MAE. **(b)** Hold-out set experimental ground-truth normalized transfection vs. optimized LGBM model-predicted normalized transfection across all five cross validation folds (n = 1080). Model predictions mirrored experimental results. **(c)** Training and validation error dependent on training set size of the selected LGBM model for B16F10. Convergence and plateauing of training and validation curves around 700 formulations indicate that our training dataset (1,080 formulations) exceeded the minimum necessary for robust model training. **(d)** Input features are removed from the training data until the removal of more features would lead to worse model performance. B16F10 LGBM model requires only 10 out of 14 features for optimal performance. **(e)** Comparison of refined model performance across cell types. HepG2, B16F10, and PC3 cell types achieved average error around 5% across five repeated 5-fold splits (n = 5).

**Figure 4.**
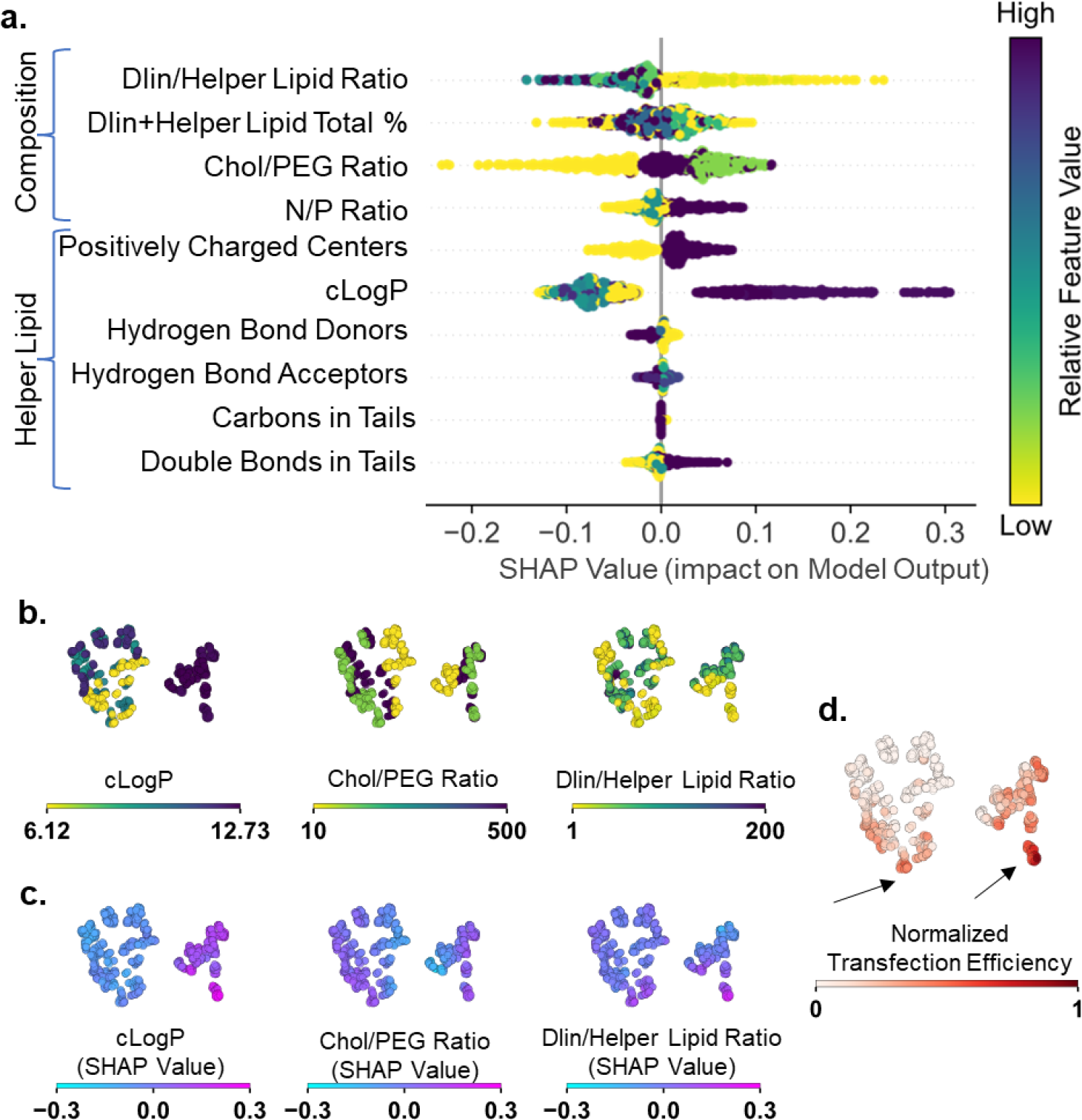
SHAP Elucidates LNP Structure-Function Relationships for B16F10 transfection. **(a)** SHAPley Additive Explanations summary plot describing the impact of feature values on final model output for the refined B16F10 LGBM model. Features are found on the y-axis, grouped by feature type (composition or helper lipid). Each point represents a single LNP formulation, and the color represents the relative feature value within the formulation. Points located to the right of the x-axis (x>0) indicate positive impact on predicted transfection, while points located to the left (x<0) indicate negative impact. **(b–d)** LNP formulations represented by individual dots embedded on 2 dimensions by t-SNE dimensional reduction of all SHAP values. **(b)** Formulations are colored by feature value used. **(c)** Formulations are colored by feature SHAP value. **(d)** Formulations colored by ground-truth normalized transfection efficiency in B16F10 cells. Black arrows indicate potential regions for improved LNP transfection efficiency.

**Figure 5.**
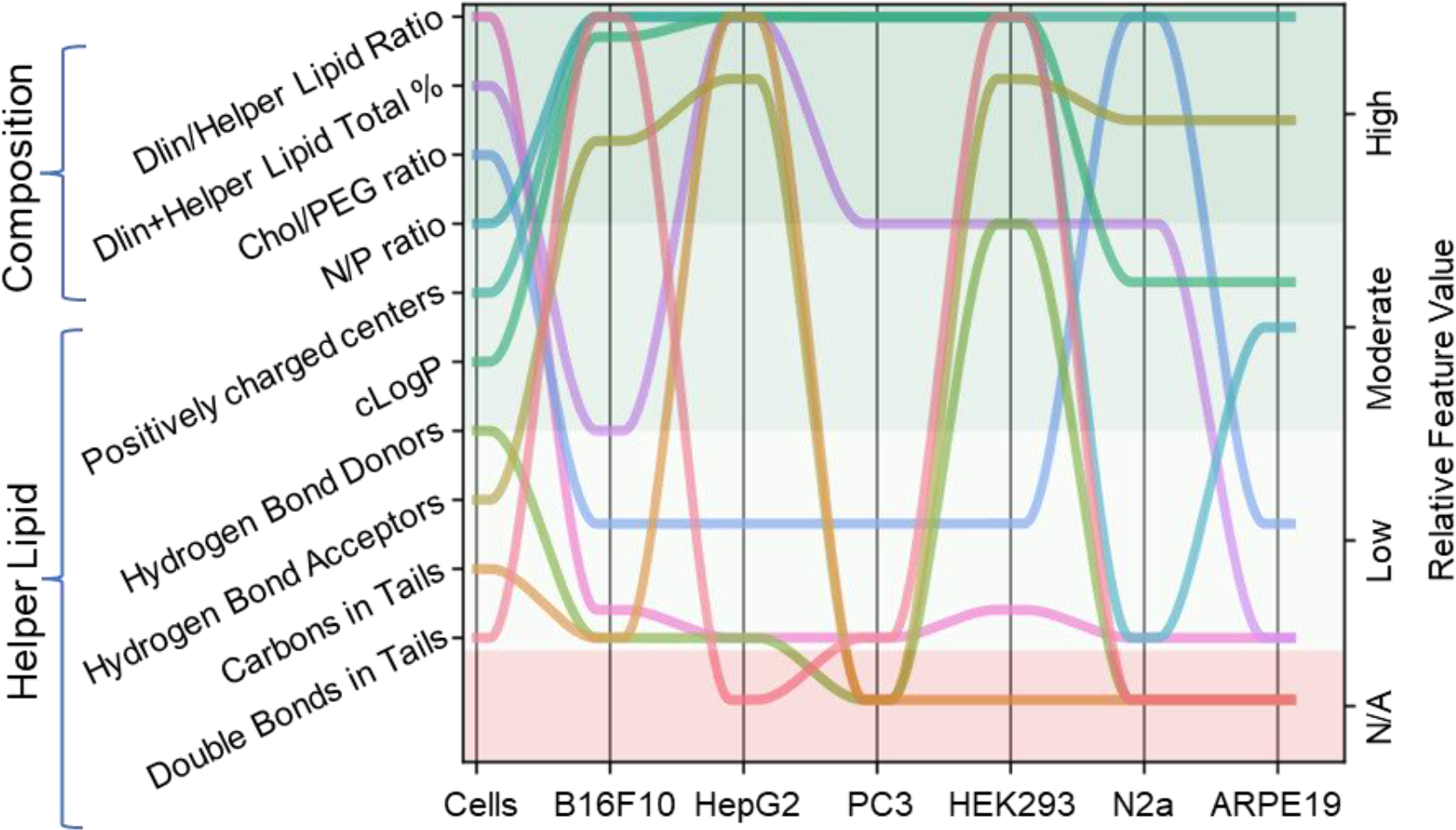
ML suggested design features of LNPs for cell-type specific transfection. Bump plot of input features (colors) and suggested LNP parameter values (y-axis) for optimal cell-type (x-axis) transfection based on SHAP values from optimized and feature refined models for each cell type. Features that are in the red “N/A” regions were not contained in the refined input features for that cell type. Differences in optimal parameter values between cell types further highlight that LNP formulation parameters must be optimized for cell type-preferential transfection.

We attribute our high model accuracies to our careful model selection process and the integration of HTS data generation ensuring the availability of reliable and well-curated datasets that focused primarily on formulation composition parameters. To validate whether the training dataset size was sufficiently large for model training, the best performing model for each cell type was retrained with a varying number of randomly selected datapoints, and the validation loss (MAE when predicting hold-out data) and the training loss (MAE when predicting training data) were recorded. In B16F10 cells, the validation loss approached the training loss; and an overall plateau of the loss curves was observed, indicating that the size of the training dataset comfortably exceeded the minimum needed for model training and validation, and that minimal overfitting was observed (**Figure 3c**). Similar trends were observed in the other cell types (**Supplemental Figure 6**). These results highlight the benefit of integrating HTS data generation in our computational analysis. Our approach also provides a major improvement for downstream mechanistic analysis compared to many other studies within the LNP field^21,22,24^. We not only ensured that the training data gathered was sufficient to construct robust and reliable models, but also minimized lab-to-lab variation and experimental reproducibility issues because all data is gathered under the same test conditions and with the same protocols. The consistency and comparability of our dataset, combined with our focus on data diversity (ensuring that low, moderate, and high performing formulations were well-represented) sets this study apart from previous work. Previously reported studies, in contrast, relied on training models with relatively small, non-curated datasets mined from historical data or literature, which often times used inconsistent protocols and narrowly focused on “optimized” results, therefore provided little understanding of why most LNP formulations fail^22,41^. In contrast, our approach leverages a larger and reliable dataset that is equally inclusive of both optimal and non-optimal formulations, enabling us to train trustworthy models that capture trends governing LNP transfection performance.

### Feature refinement establishes that chemical design features of lipid components are more predictive of transfection than nanoparticle-level design features

We next conducted feature refinement aiming to remove unnecessary features and mitigate the risk of mistakenly deriving spurious relationships between certain features. By comparing the Ward linkage distances from unsupervised hierarchical clustering using the farthest neighbor algorithm (**Supplementary Figure 7**), highly clustered features were removed from the model inputs individually. If the model performance worsened, the feature was restored^33^. Feature refinement for the B16F10 cell type showed that the model only required 10 of the initial 14 features (**Figure 3d)**. This process was repeated for each cell line individually. Impressively, our models fit using refined feature sets for B16F10 and PC3 achieved around 5% error on average, and models for all cell types achieved less than 9% error on average (**Figure 3e, Table 2**). From the initial 14 features, models for B16F10, HepG2, and HEK293 retained between 9 and 10 features, while models for N2a, PC3, and ARPE-19 retained only 7 features (**Table 2, Supplementary Figure 8**). No matter the cell type, all four formulation composition features were retained (**Table 2**). Of the helper lipid chemical features, the number of positively charged centers and cLogP was retained across all cell types; other helper lipid features such as number of hydrogen bond donors/acceptors, double bonds, and total carbon atoms in the lipid tails were necessary for models to perform optimally, but different features were retained in each cell type (**Table 2**). The number of positively charged centers contributes to the overall charge of LNPs; and cationic/zwitterionic lipids were identified more frequently in high transfecting formulations in all cell types (**Figure 2a**). The number of hydrogen bond donors/acceptors also characterizes potential hydrogen bond interactions between lipid components and the nucleic acid payload. Non-electrostatic helper lipid features helped the model differentiate between similarly charged helper lipids, particularly characterizing the structure and chemical properties of lipid tails rather than headgroup, emphasizing the importance of holistically considering all chemical properties within LNP formulations. Notably, all nanoparticle-level features such as size, PDI, and _ζ_-potential were consistently removed, and the exclusion of these features almost always improved model performance (**Supplementary Figure 8**). It is possible that some nanoparticle-level features introduced potentially spurious contributions to predictions in these cell types. This result may have been influenced by the biased distribution of LNP sizes (the majority within the 100–300 nm range) and _ζ_-potential values (the majority within -10 to +10 mV) (**Supplementary Figure 1c**). As such, there simply was not enough variability in these features for these nanoparticle-level features to contribute to transfection efficiency trends. This conclusion should also be considered within the context of LNP uptake mechanism. Most of the cell types studied favor 100–300-nm and neutral or slightly positive charged particles^42^. Since the majority of our LNP formulations fall within these size and _ζ_-potential ranges, the tested models do not need these nanoparticle-level features to make accurate predictions. Overall, the feature refinement process enabled the ML pipeline to retain the most critical input parameters and further improved performance, highlighting the importance of LNP composition and helper lipid choice. Additionally, this process will aid model interpretation and derivation of structure-function relationships during post-hoc SHAP analysis.

### Feature importance illuminates cell type-preferential LNP formulation design rules

The SHAP summary plot for the B16F10 cell type mapped the feature importance and illustrates structure-function relationships unveiled by the LGBM model and identifies relative feature values associated with enhanced or diminished transfection efficiency (**Figure 4a**). Composition parameters were strongly predictive of transfection efficiency. DLin/helper lipid ratios approaching 1:1 or 10:1, Chol/PEG ratios approaching 100:1 or 500:1, and N/P ratios approaching 12 resulted in strongly positive contributions to transfection efficiency. We did not observe obvious trends for the DLin and helper lipid total molar percentage, suggesting complex interactions between lipid components. Particularly, lower Dlin and helper lipid total molar percentage around 20–40% and higher Chol:PEG ratios near 100–500 tend to enhance their respective positive influence on transfection efficiency (**Supplementary Figure 9**). Helper lipid cLogP near 12.7–12.9, positive charges (cationic or zwitterionic lipid head groups), and greater unsaturation of helper lipid tails (2 double bonds) improve transfection in B16F10 cells (**Figure 4a**).

Visualizing the location of feature values and SHAP values of the top 3 important features in 2-dimensional space by t-distributed stochastic neighbors embedding dimensional reduction provides deeper insight into formulation design rules (**Figure 4b–c, Supplemental Figure 10**). There are two separate local or potentially global transfection maxima that can be identified within the formulation parameter space screened for B16F10 cells (**Figure 4d**). By identifying formulations residing in regions of high transfection efficiency, we discovered LNP design rules from the refined LGBM model. Helper lipid cLogP (near 12.7–12.9), Chol/PEG ratios approaching 100:1, and DLin/helper lipid ratios near 1:1 and 10:1 were all clearly localized with regions of high transfection **(Figure 4b)**. Similarly, high SHAP values colocalized in regions of high transfection and display the additive effect of optimizing each formulation parameter to improve transfection (**Figure 4c**). Other composition and helper lipid features values also show colocalization with regions of high transfection efficiency, albeit less significantly, further emphasizing the complexity of LNP formulation design (**Supplementary Figure 11, 12**). The open regions in the 2D plots (highlighted with arrows in **Figure 4d**), particularly where high values of experimental transfection efficiencies and SHAP values are localized, identifies feature spaces where improved LNP formulations may be discovered (**Figure 4d**). In future LNP libraries, we will prioritize designing formulations with helper lipid cLogP near or higher than 12.7–12.9, DLin to helper lipid ratios between 1:1 and 10:1, and Chol/PEG ratios between 100:1 and 500:1 to augment transfection performance.

SHAP feature importance analysis was repeated for all six cell types to identify LNP formulation design rules for cell type-preferential transfection (**Supplementary Figure 13**). Each cell type requires distinct and mutually exclusive sets of different formulation parameters to optimize transfection (**Table 3** and **Figure 5**). We attribute this result to the unique uptake and intracellular trafficking characteristics of each cell type. SHAP lends credence to our hypothesis that cell type-preferential transfection *in vitro* can be accomplished by modulating the formulation parameters. Notably, cell types share multiple overlapping composition features values. Most prominently, helper lipids containing cationic or zwitterionic helper lipids (1 positively charged center) and low DLin/helper lipid ratios (1:1 and 10:1) were associated with optimal transfection in all cell types. It is essential to optimize the DLin/helper lipid ratio, highlighting that even subtle alterations to formulation recipes can have far-reaching effects on transfection outcomes. SHAP analysis is effective in uncovering LNP design parameters preferred by each cell type and provides a promising approach for ML-guided design of bespoke LNPs tailored to the unique requirements of each cell type.

## Conclusion

LNP screening has primarily focused on discovering and validating new ionizable lipid chemistries while marginalizing complex interactions between lipid components (helped lipids, ionizable lipids, PEGylated lipids, and sterols). We developed a machine learning pipeline to exploit large well-curated datasets from HTS and elucidate important structure-function relationships of different LNP compositions. We trained, optimized, and validated machine learning models across multiple cell types. After model selection and hyperparameter optimization, we identified LNP design features and formulation parameters that exerted the greatest influence over transfection efficiency. SHAP analysis elucidated the physicochemical drivers of high-performing LNP formulations for each cell type. Importantly, our comparative analysis demonstrates that we must delicately modulate helper lipid chemical parameters and formulation composition to achieve cell type-preferential transfection.

While this study primarily focuses on structure-function relationships driving pDNA LNPs delivery performance, particularly helper lipid chemical identity and LNP formulation parameters, our approach is generalizable to a larger chemical design space. Our pipeline is now ideally positioned to investigate expanded parameters spaces opened up by new lipid chemistries, the addition of more lipid components (such as SORT lipids), and new payload types including mRNA and other RNA therapeutics^43^. The inclusion of additional input features characterizing the LNP formulation such as encapsulation efficiency, payload distribution, or protein corona composition would potentially improve the model’s understanding of the LNP design rules. Future work will also parametrize the unique biological signatures of target cell types, which will serve as additional input features for integrated models. This will allow us to analyze multiple cell types together within a single unified model instead of developing separate models for each cell type. Unified models will identify the governing characteristics of the target cell influencing the efficacy of LNP gene delivery (e.g., GAG expression, endocytic preferences etc.). Finally, while the current study has focused on *in vitro* LNP transfection, our pipeline can potentially be improved by incorporating alternative high-throughput *in vivo* screening techniques such as LNP barcoding ^44^ and cluster-mode screening^19^ will potentially enable researchers to uncover the mechanisms driving *in vivo* LNP structure-function relationships.

## Methods

### Preparation and characterization of pDNA LNPs

The LNPs were formulated in 96-well U-bottom plates according to a previously described method^19^, using DLin-MC3-DMA (MedKoo Biosciences), cholesterol (Sigma-Aldrich), DMG-PEG2000 (NOF America Corporation), and one of the helper lipids, DSPC, DOPE, DOTAP, DDAB, 18PG, and 14PA (all from Avanti Polar Lipids). The lipid phase was prepared by dissolving lipid components in ethanol at a predetermined concentration (based on the formulation composition); and the firefly luciferase pDNA (Aldevron) was diluted in 25 mM magnesium acetate buffer (pH 4.0). LNPs were formulated by combining the aqueous and ethanol phase at a 3:1 ratio with pipette mixing without further purification. LNPs were prepared fresh for each experiment. The size (z-avg), polydispersity index (PDI), and _ζ_-potential of the LNP library were measured via dynamic light scattering using a Nano ZS90 Zetasizer (Malvern Analytical). Aliquots of 100 μL of LNPs (25 μg pDNA/mL) were used for measuring size and PDI. For _ζ_-potential, 100 μL of LNPs were diluted into 900 μL of PBS prior to measurement in a capillary cuvette. Measurements were performed in triplicate for each test sample.

### In vitro transfection and luciferase assay

For monolayer culture studies, HepG2 cells (ATCC HB-8065), HEK293T (ATCC CRL-3216), B16F10 cells (ATCC CRL-6475), PC3 (ATCC CRL-1435), Neuro2a (ATCC CCL-131) and ARPE-19 cells (ATCC CRL-2302) cells (American Type Culture Collection, USA) were seeded into 96-well plates at a cell density of 10,000 cells per well one day prior to transfection. After formulation, LNPs were pipetted into culture medium at a final particle concentration of 1 μg pDNA/mL (i.e., 4 μL of a particle suspension at 25 μg pDNA/mL was pipetted into 100 μL of culture media in the wells. After 48 h of incubation, cells were lysed and luciferase expression was analyzed using a luminometer following a standard protocol.

### Helper lipid parameterization

The chemical features used to parameterize the helper lipid structure were carefully selected based on historical knowledge of important lipid structures. Other more complex parameterization methods such as Extended connectivity fingerprints^28^ were not used to preserve interpretability of the chemical features and due to the relatively small diversity of helper lipids tested (six in total). Helper lipid chemical features such as the number of positively and charged centers, the numbers of hydrogen bond donors and acceptors, and total carbons and double bonds in the lipid tails were included; and the calculated log of the partition coefficient between octanol and water (cLogP) and calculated topological polar surface area (cTPSA) property features were calculated using ChemDraw (v22.2.0, PerkinElmer).

### Data Preprocessing

To facilitate accurate analysis and interpretation of the experimental data, natural logarithm (ln) transformation of the luciferase assay results, serving as a target output variable, was carried out to rescale the experimental transfection efficiency. Ln transformed RLU values were then floored to the average value of blank wells (around e^2^ RLU) to minimize noise at low and near zero transfection measurements. RLU measurements were then normalized using linear scaling from 0 to 1 as a fraction of the observed maximum transfection values for the corresponding cell type dataset using Eqn. 1. as follows:

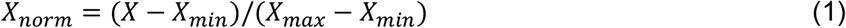

The data normalization method was applied to each cell type dataset individually to minimize the impact of cell type “transfectability” in downstream analysis.

### Model development and evaluation

The sci-kit learn python ML package (1.2.2) was used to conduct model training and prediction. The panel of models for evaluation included Multiple linear regression (MLR), Lasso regression (lasso), Partial Least Squares regression (PLS), k-Nearest Neighbors (kNN), Decision Trees (DT), Random Forest (RF), XGboost (XGB), and LightGBM (LGBM) were implemented from the publicly available sci-kit learn (v1.2.1), xgboost (v1.7.1), lightgbm (v3.3.5) python packages. A nested cross-validation procedure for training and evaluating predictive models was adapted from Aspuru-Guzik group’s GitHub repository^45^. Edits to hyperparameter grids, data splitting functions, and result saving were made for effective integration into our pipeline. This nested approach ensures robust model evaluation, hyperparameter tuning, and generalization, leading to reliable performance estimates while mitigating overfitting risks. The outer loop executed over 5-fold cross-validation splits randomly divides the data into training (80% of all data) and test sets (20% of all data). Within each outer loop iteration, an inner loop conducts a 4-fold split of the training data and runs 100 iterations of hyperparameter optimization for each split to minimize validation set mean absolute error (MAE) using a randomized grid search strategy. MAE was used to evaluate model performance because it quantifies the model‘s overall prediction error. By minimizing the MAE, we selected models that achieve a closer alignment between predictions and ground truth values, showcasing the model‘s ability to make accurate forecasts on average. The equation for the Absolute error (AE) and MAE is defined as follows:

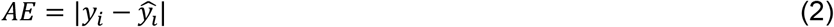

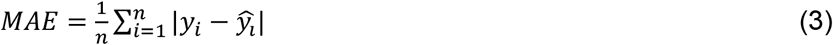

where, *n* is the number of samples, y_i_ is the predicted output by the model, and ŷ_i_is the ground-truth output.

The model hyperparameters with the lowest validation MAE from each inner loop splits was then evaluated on the outer loop hold-out test set, generating test set performance metrics including MAE, Spearman‘s rank correlation, and Pearson‘s correlation coefficient. This process was repeated for each of the five outer loop iterations. To select the final model hyperparameters, the absolute difference between inner loop validation MAE and outer loop hold-out MAE was calculated as the “score difference” for each outer-loop iteration. The model architecture with the minimum “score difference” was selected as the final model to minimize impacts of overfitting.

The model learning curve was produced by evaluating MAE as a function of the number of training datapoints provided. Different sized subsets of the training data were randomly selected and the resulting train and validation error (MAE) was averaged over 5 repeated 5-fold cross validation test-train data splits. The training size ranged from 1 to 864 formulations for the 1080-formulation datasets, and 1 to 576 for the 720-formulation datasets.

### Model refinement and interpretation

The hierarchical clustering package from SciPy (v1.10.0) was used to cluster all initial input features based on the farthest neighbor clustering algorithm. Feature reduction was performed by sequentially removing a single feature based on their Ward linkage distance, and then retraining models on the training data with reduced input parameters for each cell type. The code used was adapted from the Aspuru-Guzik Group’s Github repository^45^ and the process was adjusted such that features that negatively impacted performance were not removed from the feature set. The performance of each model following removal of select features was evaluated using average MAE across shuffled 5-fold cross-validation data splits to validate the impact of feature removal on model performance. If feature removal negatively impacted model performance (increased MAE), the feature was retained, and feature refinement continued until the removal of all features was completed. The optimal refined feature set was determined as minimum necessary features to maintain or improve the initial MAE (MAE of model trained on all input features).

Feature importance analysis was conducted using the publicly available Shapley Additive Explanations package (v0.42.1). SHAP analysis plots visualized trends between input features and model outputs. LNP design rule analysis was conducted using the calculated SHAP values, measurements of feature value impacts on the final model output, to identify feature values ranges with high average SHAP values (high average positive impact on model output). These feature ranges provide potential guidelines for the necessary input feature values to maximize model output (transfection efficiency).

## Acknowledgements

This study was partially supported by the National Institute of Allergy and Infectious Diseases (NIAID) grant U01AI155313 to H.-Q.M. and the National Institute of Biomedical Imaging and Bioengineering grant R21 EB034464 to R.K.

## Author contributions

L.C. conceptualized this study. L.C., A.A., E.T., R.K. and H.-Q.M. formed the study design and developed the major conclusions. L.C., A.A., E.T., and C.S. developed the machine learning pipeline. Y.Z., J.M., E.T., W.S. and G.W. performed formulation experiments and data collection for LNP transfection performance and physical characterizations of the LNP formulations. Y.Z., J.M., A.A, E.T. contributed to data analyses and scientific discussions. H.-Q.M. and R.K. obtained funding and supervised the study. The manuscript was written by L.C., A.A., E.T., C.S., R.K. and H.-Q.M., and all other authors contributed to the review and revision of the manuscript. R.K. and H.-Q.M. contributed equally to this work as joint corresponding authors.

## Ethics declarations

### Reporting summary

Further information on research design is available in the Nature Research Reporting Summary linked to this article.

## Data availability

All data needed to evaluate the conclusions in this paper are present in the paper and/or the Supplementary Materials. Source data are provided with this paper.

## Code availability

The code in python for the data preprocessing, model selection, feature reduction, and SHAP analysis are available from the authors upon request.

## References

1. Bulaklak, K. & Gersbach, C. A. The once and future gene therapy. Nat. Commun. 11, 5820 (2020).

2. Sung, Y. & Kim, S. Recent advances in the development of gene delivery systems. Biomater. Res. 23, 8 (2019).

3. Wang, D., Tai, P. W. L. & Gao, G. Adeno-associated virus vector as a platform for gene therapy delivery. Nat. Rev. Drug Discov. 18, 358–378 (2019).

4. Ren, D., Fisson, S., Dalkara, D. & Ail, D. Immune responses to gene editing by viral and non-viral delivery vectors used in retinal gene therapy. Pharmaceutics 14, 1973 (2022).

5. Shirley, J. L., De Jong, Y. P., Terhorst, C. & Herzog, R. W. Immune responses to viral gene therapy vectors. Mol. Ther. 28, 709–722 (2020).

6. Tenchov, R., Bird, R., Curtze, A. E. & Zhou, Q. Lipid nanoparticles─From liposomes to mRNA vaccine delivery, a landscape of research diversity and advancement. ACS Nano 15, 16982–17015 (2021).

7. Jung, H. N., Lee, S.-Y., Lee, S., Youn, H. & Im, H.-J. Lipid nanoparticles for delivery of RNA therapeutics: Current status and the role of in vivo imaging. Theranostics 12, 7509–7531 (2022).

8. Yang, L. et al. Recent advances in lipid nanoparticles for delivery of mRNA. Pharmaceutics 14, 2682 (2022).

9. Huang, Q., Zeng, J. & Yan, J. COVID-19 mRNA vaccines. J. Genet. Genomics 48, 107–114 (2021).

10. Hald Albertsen, C. et al. The role of lipid components in lipid nanoparticles for vaccines and gene therapy. Adv. Drug Deliv. Rev. 188, 114416 (2022).

11. Sun, D. & Lu, Z.-R. Structure and function of cationic and ionizable lipids for nucleic acid delivery. Pharm. Res. 40, 27–46 (2023).

12. Aldosari, B. N., Alfagih, I. M. & Almurshedi, A. S. Lipid nanoparticles as delivery systems for RNA-based vaccines. Pharmaceutics 13, 206 (2021).

13. Patel, S. et al. Naturally-occurring cholesterol analogues in lipid nanoparticles induce polymorphic shape and enhance intracellular delivery of mRNA. Nat. Commun. 11, 983 (2020).

14. Kulkarni, J. A., Witzigmann, D., Leung, J., Tam, Y. Y. C. & Cullis, P. R. On the role of helper lipids in lipid nanoparticle formulations of siRNA. Nanoscale 11, 21733–21739 (2019).

15. Zong, Y., Lin, Y., Wei, T. & Cheng, Q. Lipid nanoparticle (LNP) enables mRNA delivery for cancer therapy. Adv. Mater. 2303261 (2023) doi:10.1002/adma.202303261.

16. Zhao, X. et al. Imidazole‐based synthetic lipidoids for in vivo mRNA delivery into primary T lymphocytes. Angew. Chem. Int. Ed. 59, 20083–20089 (2020).

17. Ni, H. et al. Piperazine-derived lipid nanoparticles deliver mRNA to immune cells in vivo. Nat. Commun. 13, 4766 (2022).

18. Guimaraes, P. P. G. et al. Ionizable lipid nanoparticles encapsulating barcoded mRNA for accelerated in vivo delivery screening. J. Controlled Release 316, 404–417 (2019).

19. Zhu, Y. et al. Multi-step screening of DNA/lipid nanoparticles and co-delivery with siRNA to enhance and prolong gene expression. Nat. Commun. 13, 4282 (2022).

20. Zhu, Y. et al. Screening for lipid nanoparticles that modulate the immune activity of helper T cells towards enhanced antitumor activity. Nat. Biomed. Eng. In Press, (2023).

21. Gong, D. et al. Machine learning guided structure function predictions enable in silico nanoparticle screening for polymeric gene delivery. Acta Biomater. 154, 349–358 (2022).

22. Wang, W. et al. Prediction of lipid nanoparticles for mRNA vaccines by the machine learning algorithm. Acta Pharm. Sin. B 12, 2950–2962 (2022).

23. Maharjan, R. et al. Comparative study of lipid nanoparticle-based mRNA vaccine bioprocess with machine learning and combinatorial artificial neural network-design of experiment approach. Int. J. Pharm. 640, 123012 (2023).

24. Gao, H. et al. Development of in silico methodology for siRNA lipid nanoparticle formulations. Chem. Eng. J. 442, 136310 (2022).

25. Tamasi, M. J. et al. Machine learning on a robotic platform for the design of polymer– protein hybrids. Adv. Mater. 34, 2201809 (2022).

26. Nohara, Y., Matsumoto, K., Soejima, H. & Nakashima, N. Explanation of machine learning models using improved Shapley additive explanation. Proc. 10th ACM Inter Conf Bioinformatics, Computational Biology Health Informatics. p. 546. Association for Computing Machinery (New York, NY, 2019).

27. Li, S. et al. Payload distribution and capacity of mRNA lipid nanoparticles. Nat. Commun. 13, 5561 (2022).

28. Rogers, D. & Hahn, M. Extended-Connectivity Fingerprints. J. Chem. Inf. Model. 50, 742–754 (2010).

29. Cheng, X. & Lee, R. J. The role of helper lipids in lipid nanoparticles (LNPs) designed for oligonucleotide delivery. Adv. Drug Deliv. Rev. 99, 129–137 (2016).

30. Mason, R. D., Lind, D. A. & Marchal, W. G. Statistics: An Introduction. 5th Ed. (Duxbury Press, 1998).

31. Carozza, G. et al. New insights into dose-dependent effects of curcumin on ARPE-19 Cells. Int. J. Mol. Sci. 23, 14771 (2022).

32. Mannino, G. et al. ARPE-19 conditioned medium promotes neural differentiation of adipose-derived mesenchymal stem cells. World J. Stem Cells 13, 1783–1796 (2021).

33. Bannigan, P. et al. Machine learning models to accelerate the design of polymeric longacting injectables. Nat. Commun. 14, 35 (2023).

34. Kotu, V. & Deshpande, B. Classification. in Data Science: Concepts and Practice. 2nd Ed. pp. 65–163 (Elsevier, 2018).

35. Nathanael, K., Cheng, S., Kovalchuk, N. M., Arcucci, R. & Simmons, M. J. H. Optimization of microfluidic synthesis of silver nanoparticles: A generic approach using machine learning. Chem. Eng. Res. Des. 193, 65–74 (2023).

36. Yakoubi, S. et al. Recent advances in delivery systems optimization using machine learning approaches. Chem. Eng. Process. - Process Intensif. 188, 109352 (2023).

37. Yu, F., Wei, C., Deng, P., Peng, T. & Hu, X. Deep exploration of random forest model boosts the interpretability of machine learning studies of complicated immune responses and lung burden of nanoparticles. Sci. Adv. 7, eabf4130 (2021).

38. Batra, R. et al. Machine learning overcomes human bias in the discovery of self-assembling peptides. Nat. Chem. 14, 1427–1435 (2022).

39. Loecher, A., Bruyns-Haylett, M., Ballester, P. J., Borros, S. & Oliva, N. A machine learning approach to predict cellular uptake of pBAE polyplexes. Biomater. Sci. 11, 5797–5808 (2023).

40. Yamankurt, G. et al. Exploration of the nanomedicine-design space with high-throughput screening and machine learning. Nat. Biomed. Eng. 3, 318–327 (2019).

41. Metwally, A. A., Nayel, A. A. & Hathout, R. M. In silico prediction of siRNA ionizable-lipid nanoparticles in vivo efficacy: Machine learning modeling based on formulation and molecular descriptors. Front. Mol. Biosci. 9, 1042720 (2022).

42. Sousa De Almeida, M. et al. Understanding nanoparticle endocytosis to improve targeting strategies in nanomedicine. Chem. Soc. Rev. 50, 5397–5434 (2021).

43. Cheng, Q. et al. Selective organ targeting (SORT) nanoparticles for tissue-specific mRNA delivery and CRISPR–Cas gene editing. Nat. Nanotechnol. 15, 313–320 (2020).

44. Dahlman, J. E. et al. Barcoded nanoparticles for high throughput in vivo discovery of targeted therapeutics. Proc. Natl. Acad. Sci. 114, 2060–2065 (2017).

45. Aspuru-Guzik, A. long-acting-injectables. GitHub repository. (2022). At <https://github.com/aspuru-guzik-group/long-acting-injectables>

