## Supplemental Information for "Machine Learning Elucidates Design Features of Plasmid DNA Lipid Nanoparticles for Cell Type-Preferential Transfection"

**Supplementary Table 1: Formulation Composition Features Ranges**

| Composition Parameter | Parameter Values |
|---|---|
| DLin/helper lipid ratio | 1, 10, 50, 100, 200 |
| DLin + helper lipid total % | 20, 40, 60, 80 |
| Chol/PEG ratio | 10, 100, 500 |
| N/P ratio | 4, 8, 12 |

**Supplementary Table 2: Helper Lipid Chemical Parameterization**

| Parameter Name | Parameter Description | Helper Lipid | | | | | |
|---|---|---|---|---|---|---|---|
| | | DOTAP | DDAB | DOPE | DSPC | 14PA | 18PG |
| P_charged_ centers | # of positively charged centers at physiological conditions | 1 | 1 | 1 | 1 | 0 | 0 |
| N_charged_ centers | # of negatively charged centers at physiological conditions | 0 | 0 | 1 | 1 | 1 | 1 |
| cLogP | Calculated Partition coefficient between octanol and water | 12.52 | 12.73 | 9.78 | 6.12 | 11.85 | 9.30 |
| Hbond_A | # of hydrogen bond acceptors | 4 | 0 | 9 | 8 | 8 | 10 |
| Hbond_D | # of hydrogen bond donors | 0 | 0 | 2 | 0 | 2 | 3 |
| N_carbon_tails | Total # of carbon atoms in all tails | 36 | 36 | 36 | 36 | 28 | 36 |
| Double_bonds | Total # of carbon-carbon double bonds in the lipid tails | 2 | 0 | 2 | 0 | 0 | 1 |

**Supplementary Table 3. Hyperparameter Optimization Grid**

| Models | Hyperparameter Grid | Optimized Hyperparameters for B16F10 |
|---|---|---|
| LGBM | "n_estimators":[100,150,200,250,300,400,500,600], 'boosting_type': ['gbdt', 'dart', 'goss'], 'num_leaves':[16,32,64,128,256], 'max_bin': [10, 20, 30], 'max_depth': [2, 6, 10, 14, 18, 22, 26, 30], 'learning_rate':[0.1,0.01,0.001,0.0001], 'min_child_weight': [0.001,0.01,0.1,1.0,10.0], 'subsample': [0.4,0.6,0.8,1.0], 'min_child_samples':[2,10,20,40,100], 'reg_alpha': [0, 0.005, 0.01, 0.015], 'reg_lambda': [0, 0.005, 0.01, 0.015] | {'subsample': 1.0, 'reg_lambda': 0.015, 'reg_alpha': 0.005, 'num_leaves': 16, 'n_estimators': 100, 'min_child_weight': 10.0, 'min_child_samples': 10, 'max_depth': 22, 'max_bin': 10, 'learning_rate': 0.1, 'boosting_type': 'gbdt'} |
| RF | 'n_estimators':[100,300,400], 'criterion':['squared_error', 'absolute_error'], 'max_depth':[2, 6, 10, 14, 18, 22, 26, 30], 'min_samples_split':[2,4,6,8], 'min_samples_leaf':[1,2,4], 'min_weight_fraction_leaf':[0.0], 'max_features': [None, 'sqrt'], 'max_leaf_nodes':[None], 'min_impurity_decrease': [0.0], 'bootstrap':[True], 'oob_score':[True], 'ccp_alpha': [0, 0.005, 0.01] | {'oob_score': True, 'n_estimators': 400, 'min_weight_fraction_leaf': 0.0, 'min_samples_split': 2, 'min_samples_leaf': 1, 'min_impurity_decrease': 0.0, 'max_leaf_nodes': None, 'max_features': None, 'max_depth': 14, 'criterion': 'squared_error', 'ccp_alpha': 0, 'bootstrap': True} |
| XGB | 'booster': ['gbtree', 'gblinear', 'dart'], "n_estimators":[100, 150, 300, 400], 'max_depth':[3, 4, 5, 6, 7, 8, 9, 10], 'gamma':[0, 2, 4, 6, 8, 10], 'learning_rate':[0.3, 0.2, 0.1, 0.05, 0.01], 'subsample': [0.5, 0.6, 0.7, 0.8, 0.9, 1.0], 'min_child_weight': [1.0, 2.0, 4.0, 5.0], 'max_delta_step':[1, 2, 4, 6, 8, 10], 'reg_alpha':[0.001, 0.01, 0.1], 'reg_lambda': [0.001, 0.01, 0.1] | {'subsample': 0.9, 'reg_lambda': 0.01, 'reg_alpha': 0.001, 'n_estimators': 100, 'min_child_weight': 4.0, 'max_depth': 10, 'max_delta_step': 10, 'learning_rate': 0.3, 'gamma': 0, 'booster': 'dart'} |
| DT | 'criterion':['squared_error', 'friedman_mse', 'absolute_error', 'poisson'], 'splitter':['best', 'random'], 'max_depth':[None], 'min_samples_split':[2,4,6], 'min_samples_leaf':[1,2,4], 'max_features': [None, 1.0, 'sqrt','log2'], 'ccp_alpha': [0, 0.05, 0.1, 0.15] | {'splitter': 'random', 'min_samples_split': 4, 'min_samples_leaf': 2, 'max_features': None, 'max_depth': None, 'criterion': 'squared_error', 'ccp_alpha': 0} |
| kNN | 'n_neighbors':[2, 4, 5, 6, 8, 10, 12, 15, 20, 25, 30, 50], 'weights': ["uniform", 'distance'], 'algorithm': ['auto', 'ball_tree', 'kd_tree', 'brute'], 'leaf_size': [10, 30, 50, 75, 100], 'p':[1, 2], 'metric': ['minkowski'] | {'weights': 'distance', 'p': 1, 'n_neighbors': 4, 'metric': 'minkowski', 'leaf_size': 50, 'algorithm': 'auto'} |
| MLR | 'fit_intercept':[True, False], 'positive':[True, False] | {'positive': False, 'fit_intercept': False} |
| PLS | 'n_components':[2, 4, 6], 'max_iter': [250, 500, 750, 1000] | {'n_components': 6, 'max_iter': 250} |
| lasso | 'alpha':[0.01, 0.02, 0.05, 0.1, 0.25, 0.5, 1.0], 'positive':[True, False] | {'positive': False, 'alpha': 0.01} |

**Supplementary Table 4. Composition feature ranges for *in vitro* validation**

| **Formulation Composition Parameter** | **Parameter Values in Training Data** | **Parameter Values used for *in vitro* Validation** |
|---|---|---|
| DLin/helper lipid ratio | 1, 10, 50, 100, 200 | 1, 4, 5, 7, 10, 25, 30, 35, 50, 65, 75, 85, 125, 135, 150, 165, 175, 200 |
| DLin + helper lipid total % | 20, 40, 60, 80 | 20, 25, 30, 35, 40, 45, 50, 55, 60, 65, 70, 75, 80 |
| Chol/PEG ratio | 10, 100, 500 | 10, 25, 40, 50, 70, 75, 100, 200, 250, 300, 350, 400, 500 |
| N/P ratio | 4, 8, 12 | 4, 5, 6, 7, 8, 9, 10, 11,12 |

**Supplementary Table 5. Model performance during model selection and feature refinement**

| Steps Completed | Model | **HepG2** | **PC3** | **HEK293** | **B16** | **N2a** | **ARPE19** |
|---|---|---|---|---|---|---|---|
| Both | Best Model | LGBM | LGBM | LGBM | LGBM | LGBM | LGBM |
| Model Selection | MAE | 0.052 | 0.052 | 0.09 | 0.055 | 0.065 | 0.107 |
| | Spearman | 0.68 | 0.66 | 0.82 | 0.90 | 0.88 | 0.85 |
| | Pearson | 0.88 | 0.90 | 0.80 | 0.91 | 0.90 | 0.85 |
| Feature Reduction | MAE | 0.049 | 0.051 | 0.083 | 0.049 | 0.058 | 0.093 |
| | Spearman | 0.70 | 0.67 | 0.85 | 0.91 | 0.90 | 0.88 |
| | Pearson | 0.88 | 0.90 | 0.83 | 0.93 | 0.92 | 0.89 |

**Supplementary Figures**

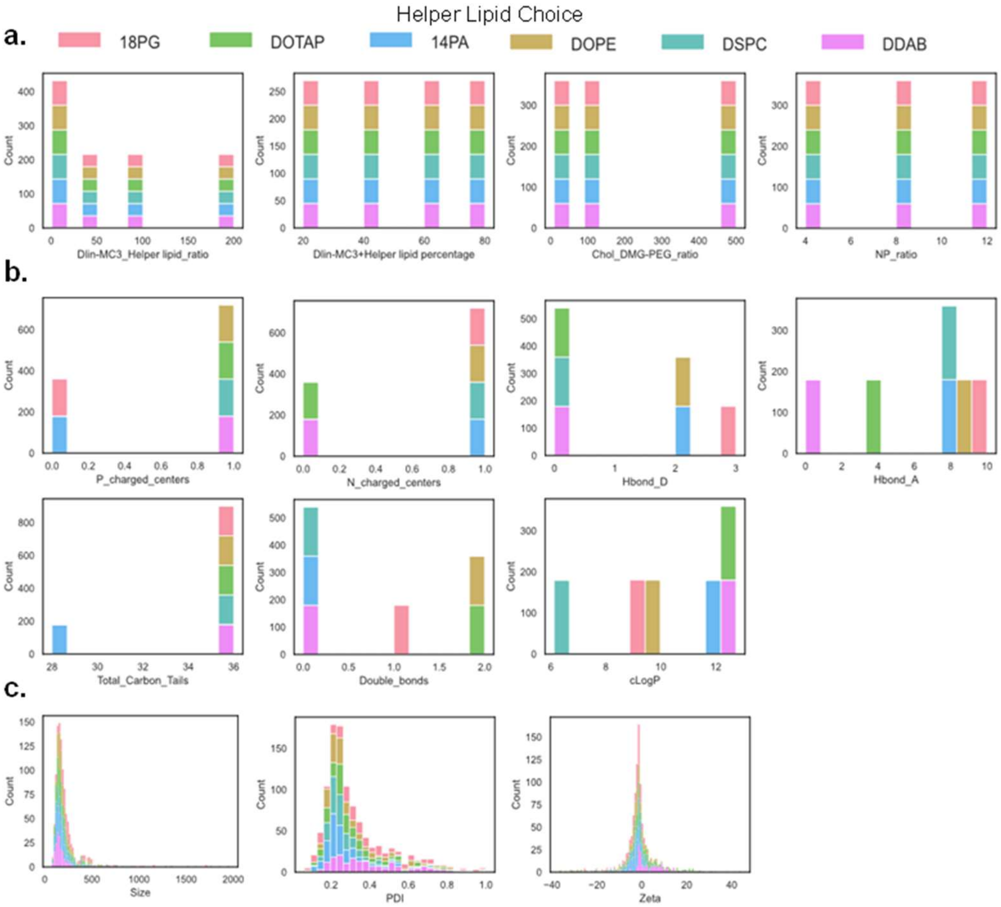


**Supplementary Figure 1: Input Parameter Distributions in 1080 formulation LNP Library.** Nanoparticle-level features, sizes, and zeta potential values of LNPs did not vary much across our LNP library. In contrast, helper lipid chemical descriptors and formulation parameters spanned a broad range of values.

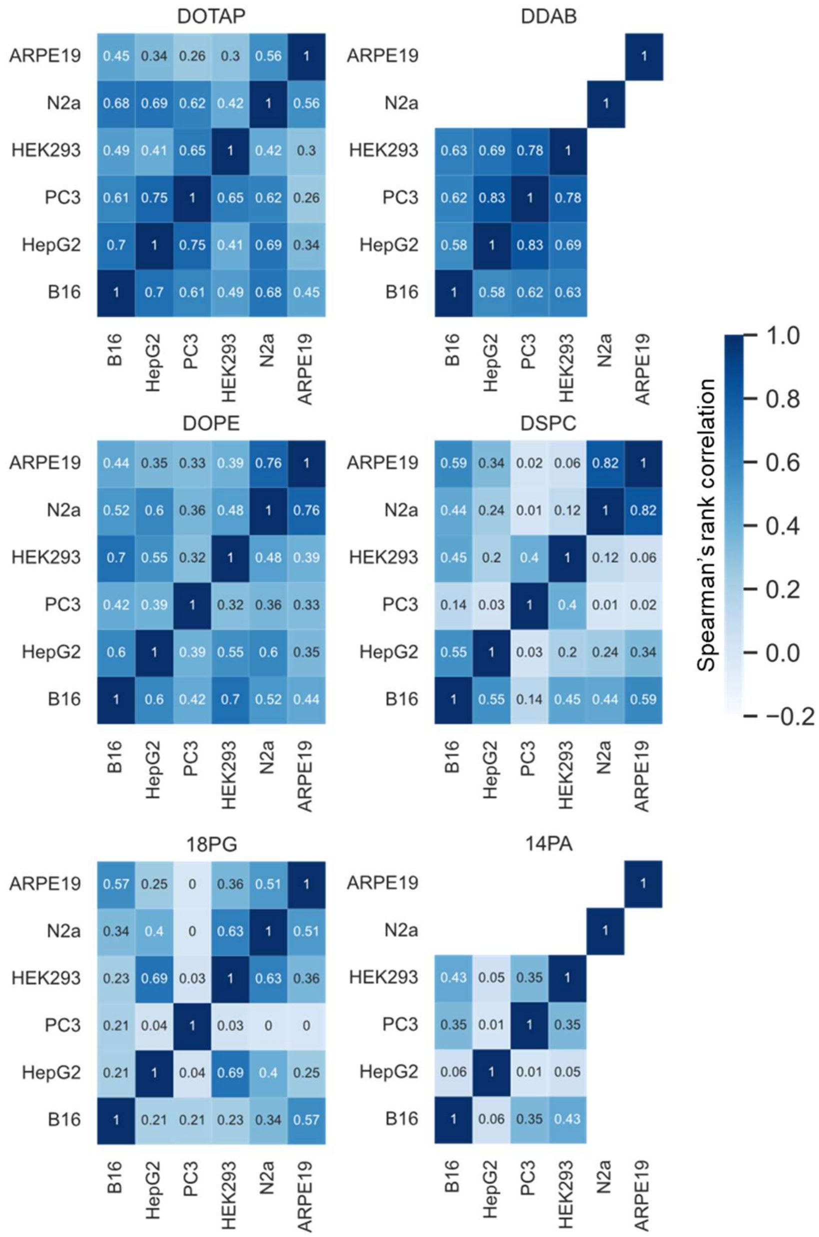


**Supplementary Figure 2. Heatmap of Transfection Correlations by Helper Lipid.** Higher values of correlation coefficients indicate overlap in transfection-driving LNP design features between two cell types. Lower values, on the other hand, suggest that LNP design criteria diverge among two cell types. For instance, N2a and ARPE19 had the highest correlation coefficients, indicating that LNPs developed for N2a are likely to perform well in ARPE-19.

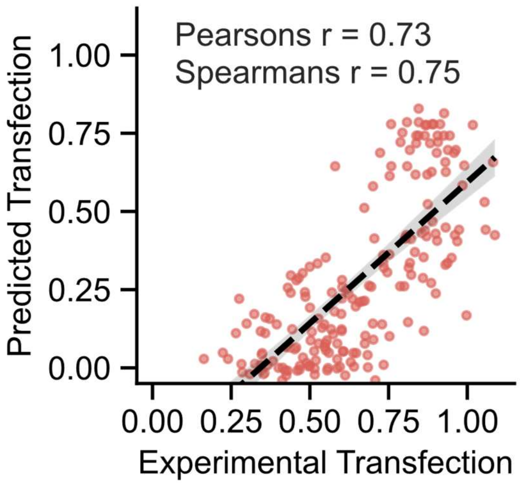


**Supplemental Figure 3*. In vitro* validation of refined LGBM model for the B16F10 cell type** (*n* = 192). Transfection is normalized to the training dataset. Strong correlation between ground-truth experimental transfection and model-predicted. Rightward shift of the linear regression suggests that the model is generally under-predicting LNP transfection efficiency. Nevertheless, the formulations with high predicted transfection efficiency still achieve high transfection *in vitro*.

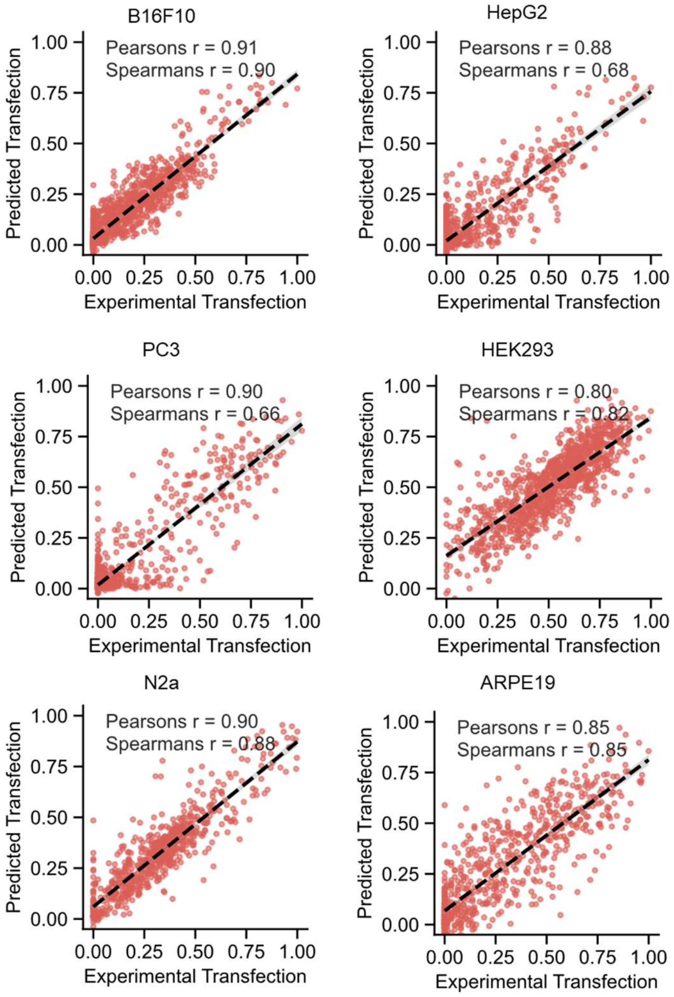


**Supplementary Figure 4. Hold-out Set Performance for all 6 Cell types.** B16F10 data has been shown in **Main Figure 3b**. It is included here for comparison purposes. Experimental results from the hold-out dataset (where data was not used for model training) closely mirrored model predictions for all six cell types, establishing the generalizability of our models (n = 1080 for B16f10, HepG2, PC3, HEP293 and n = 720 for N2a and ARPE19).

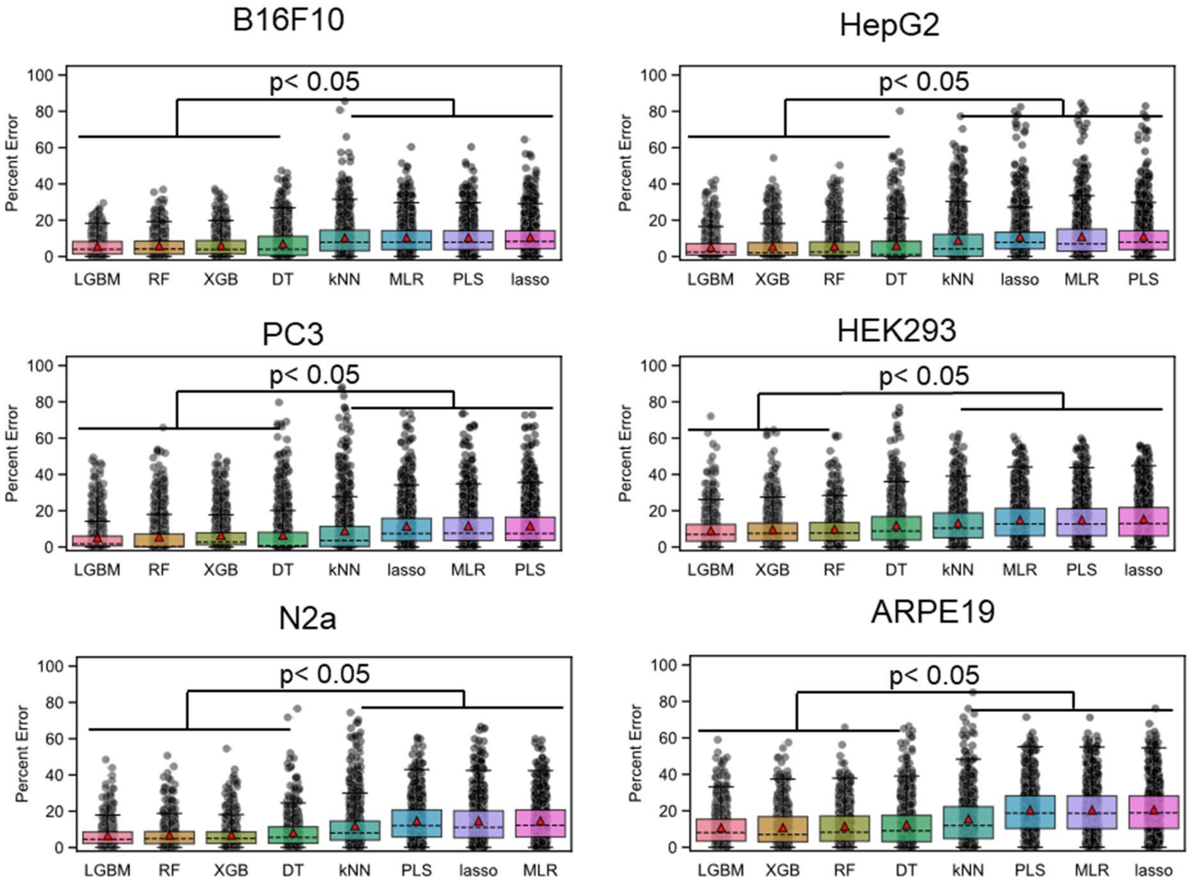


**Supplementary Figure 5. Model Selection Results for all 6 Cell types.** B16F10 data has been shown in **Main Figure 3a**. It is included here for comparison purposes. Across all cell types, decision tree-based models (LGBM, DT, RF, XGB) had lower values of MAE compared to other model architectures. Since dataset trends could be captured and described accurately by decision tree-based models alone, it indicates that our dataset is characterized by highly complex non-linear relationships between LNP features and transfection efficiency.

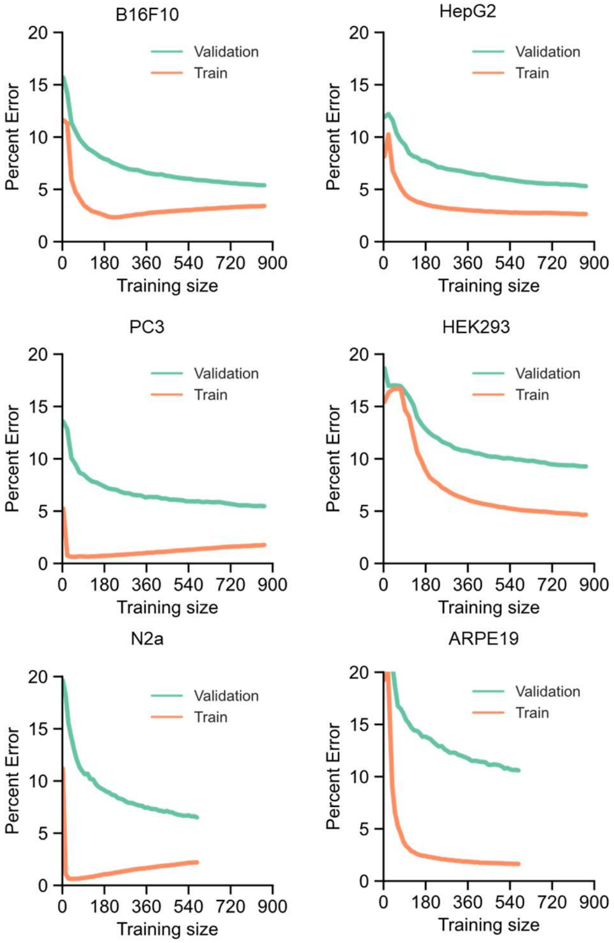


**Supplementary Figure 6. Learning Curve for all 6 Cell types.** B16F10 data has been shown in **Main Figure 3c**. It is included here for comparison purposes. For all cell types, validation error approached training error once the dataset size reached 500–800 data points. Our training dataset (1080 or 720 LNP formulations) comfortably exceeds the minimum dataset size needed to develop a generalizable model.

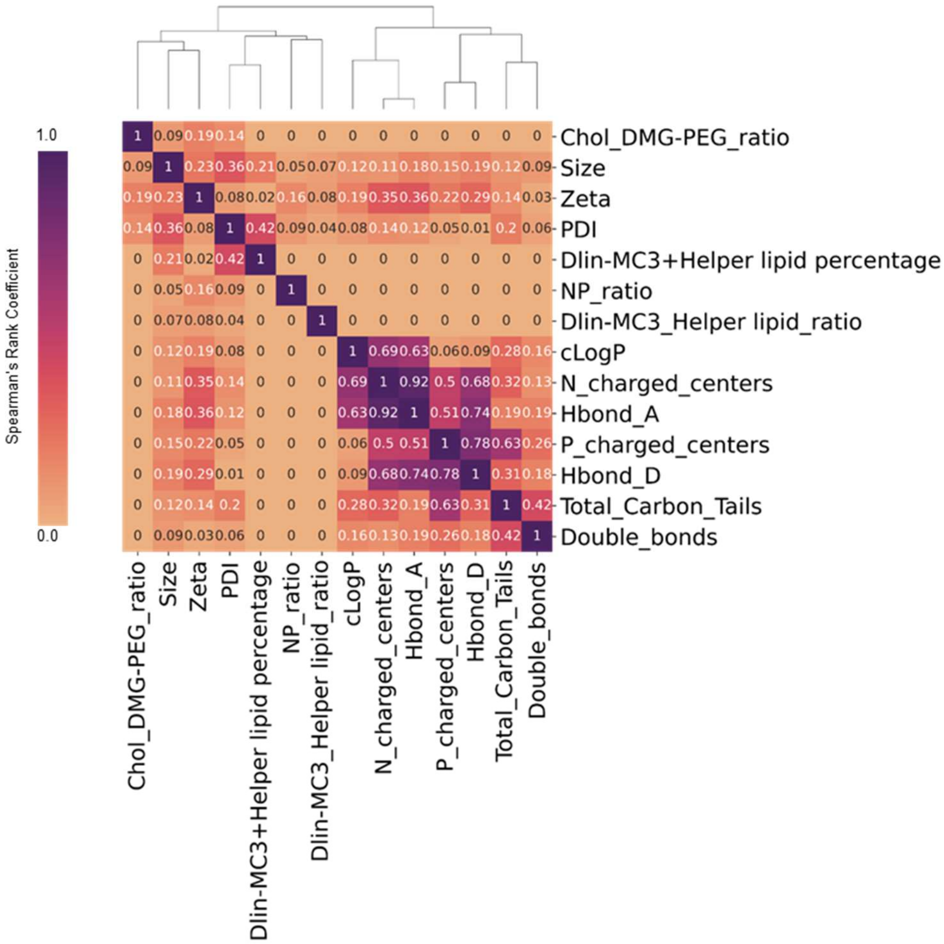


**Supplementary Figure 7. Input feature correlation heatmap and clustering**. **(a)** Heatmap of the absolute spearman's rank correlation coefficient between input feature values of the 1080-formulation dataset. **(b)** Dendrogram displays feature similarities by ward-linkage distances from farthest neighbor hierarchical clustering approach. Dendrogram representation reveals feature hierarchy by highlighting highly correlated feature pairs (with high correlation coefficients) and independent features (correlation coefficients of zero).

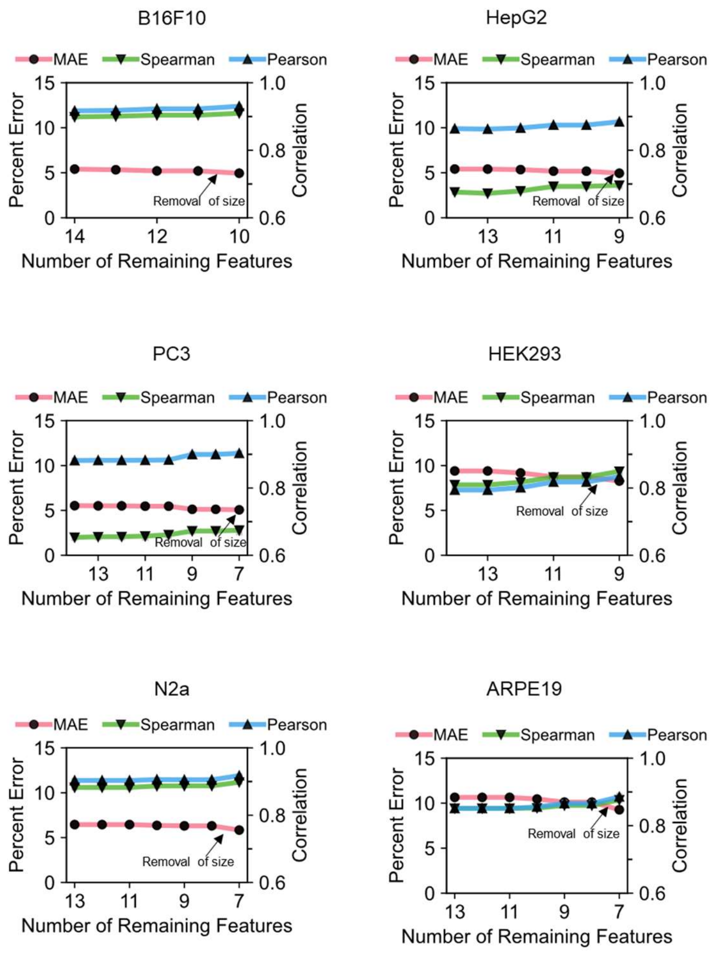


**Supplementary Figure 8. Feature Reduction for all 6 cell types.** B16F10 data has been shown in **Main Figure 3c**. It is included here for comparison purposes. The predictive value of each LNP feature is highly dependent on cell type; some cell types needed more feature refinement (needing only 7 features out of 14 to achieve high model accuracy) while others did not (retaining 9 out of 14 features during model refinement). Feature refinement for N2a and ARPE19 starts with 13 total features because the total number of carbons in the lipid tails feature does not change in value among the four helper lipids (DOTAP, DOPE, DSPC, and 18PG) screened.

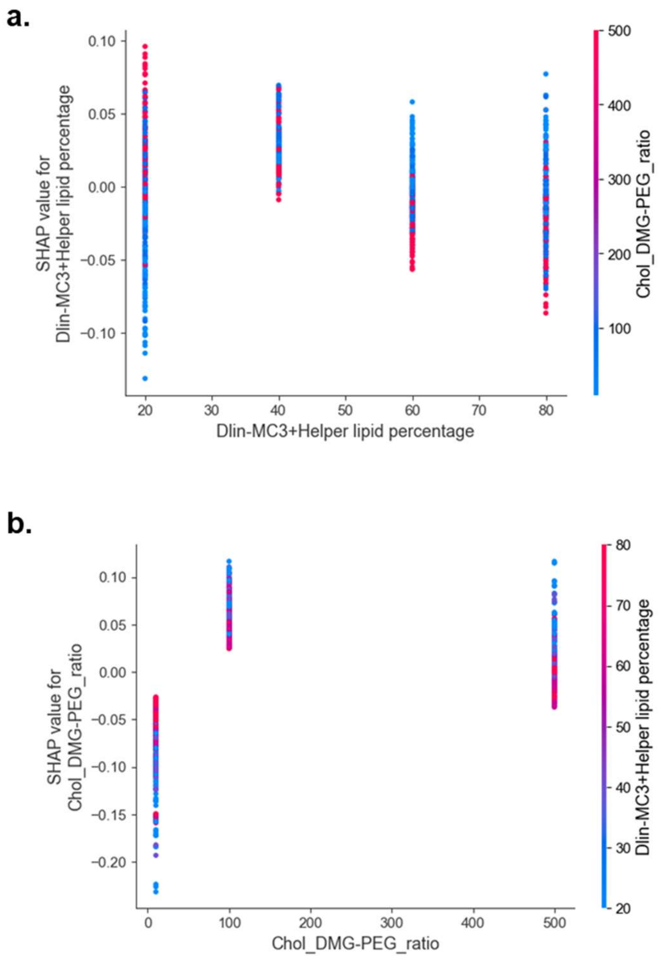


**Supplementary Figure 9. SHAP dependence plot for DLin/helper lipid ratio and Chol/PEG ratio SHAP values for the B16F10 LGBM model. (a)** Shows all 1080 formulations (dots) shifted horizontally based on DLin/Helper Lipid total percentage and colored by Chol/PEG ratio. Localization of red dots in positive SHAP values and blue in negative SHAP values show positive interaction between high Chol/PEG ratio and DLin/Helper Lipid total percentage. **(b)** Displays the same formulations with Chol/PEG ratio (X-axis) and DLin/Helper lipid percentage (colors). Low DLin/Helper lipid total percentage (blue dots) further increase SHAP values at 100 and 500 Chol/PEG ratio, but further decrease at 10 Chol/PEG ratio. This result may stem from balancing the total amount PEG lipid in the formulation.

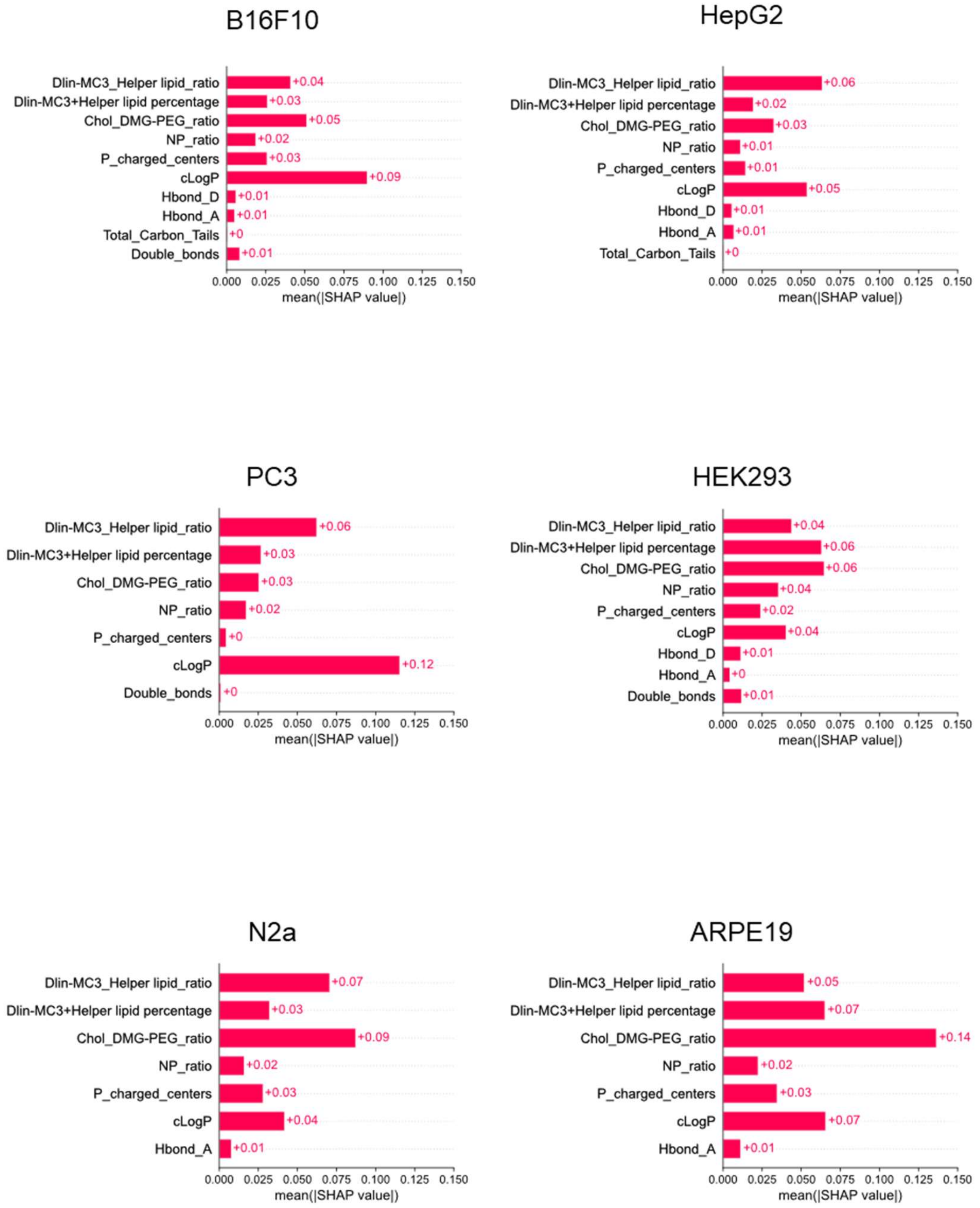


**Supplementary Figure 10. SHAP feature importance plots for all 6 cell types.** Feature importance values of helper lipid chemical descriptors and LNP formulation parameters varied widely across cell types, indicating that LNP feature optimization is highly cell type dependent.

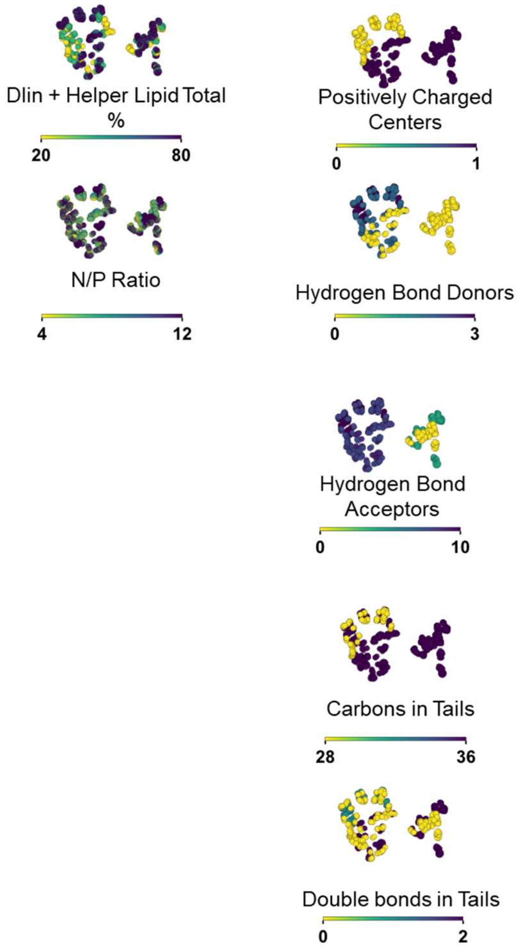


**Supplementary Figure 11. Remaining t-SNE embedded formulations colored by feature value.** LNP formulations represented by individual dots embedded on 2 dimensions by t-SNE of all SHAP values. Formulations colored by feature value used. Left side shows the remaining composition features (Chol/PEG ratio and DLin/Helper lipid ratio are shown in Main Figure 4b). Right side shows the remaining helper lipid features (cLogP is shown in Main Figure 4b). Distinct colocalization of specific feature values with regions of high transfection (Main Figure 4d) display potentially optimal feature values.

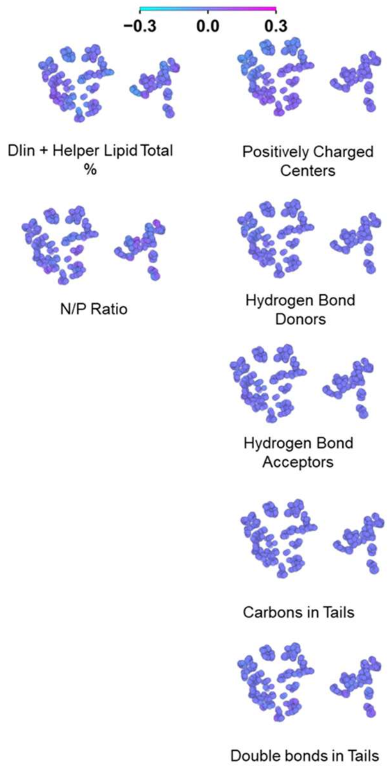


**Supplementary Figure 12. Remaining t-SNE embedded formulations colored by feature SHAP value.** LNP formulations represented by individual dots embedded on 2 dimensions by t-SNE of all SHAP values. Formulations colored by feature value used. Left side shows the remaining composition features (Chol/PEG ratio and Dlin/Helper lipid Ratio are shown in Main Figure 4c). Right side shows the remaining helper lipid features (cLogP is shown in Main Figure 4c). Colocalization of SHAP values with regions of high transfection (Main Figure 4d) display the impact of multiple interacting features on promoting LNP transfection efficiency.

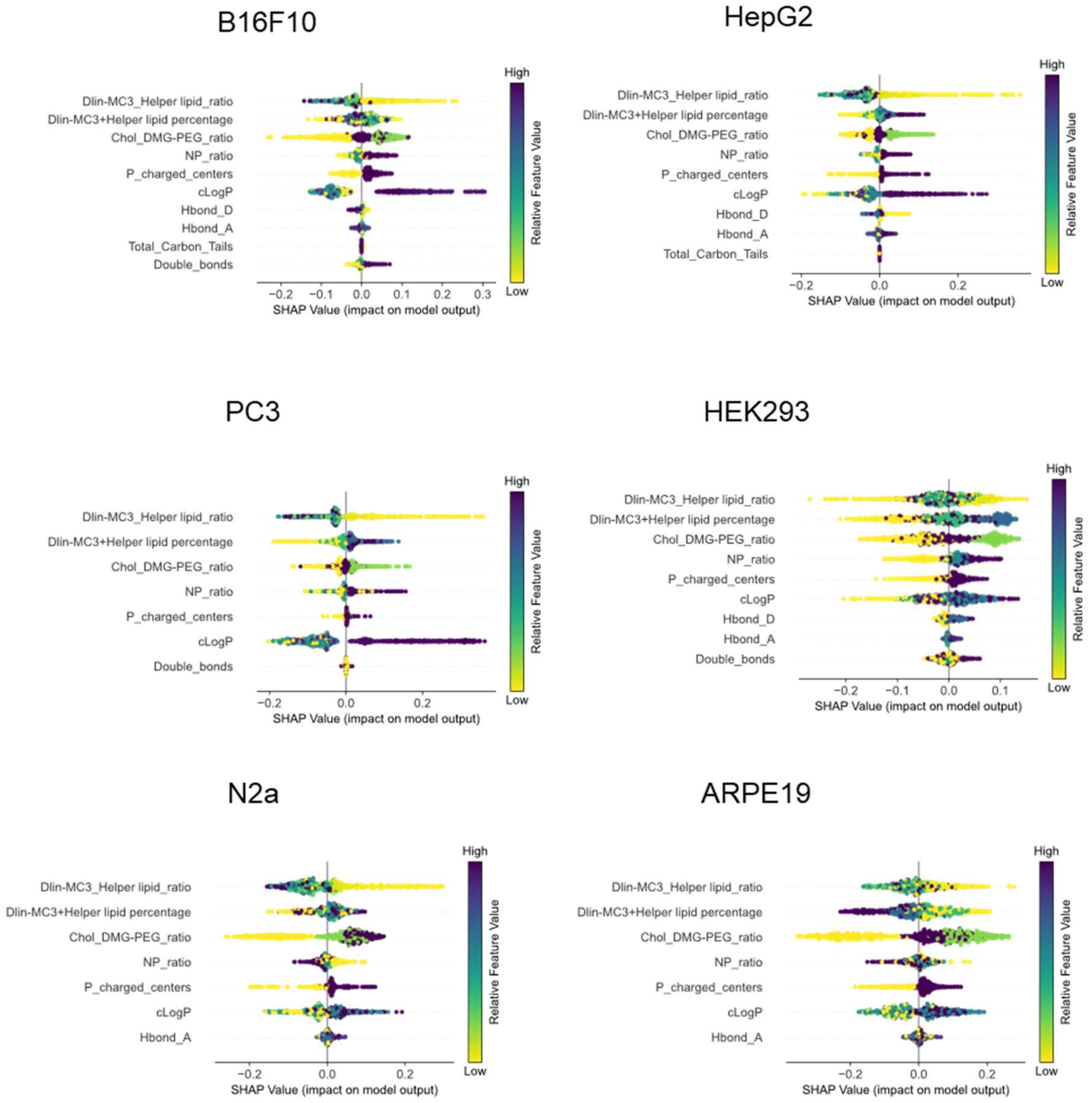


**Supplementary Figure 13. SHAP summary plots for all 6 cell types.** B16F10 data has been shown in **Main Figure 4a**. It is included here for comparison purposes. Protagonist (high feature values further right on the X-axis promote transfection) and antagonist (low feature values further right on the X-axis promote transfection) LNP features for each cell type are highlighted.
